# Camera trapping and passive acoustic monitoring as non-invasive techniques for the study of small mammals

**DOI:** 10.64898/2026.02.24.707730

**Authors:** Lili M. Stiff, Andrew Wakefield, Sean A. Rands

## Abstract

Terrestrial small mammals are of significant importance worldwide in providing valuable ecosystem services and acting as invasive species, pests, and carriers of disease. Many species are under threat from anthropogenic stressors and non-invasive techniques are required to sustain long-term and large-scale monitoring of these animals. This research compares the application of camera trapping and passive acoustic monitoring for the study of terrestrial small mammals. We conducted a field study aiming to evaluate three camera trapping tunnel designs (Modified Littlewood, Mostela and a new design utilising readily available components), and compare the performance of passive acoustic monitoring to camera trapping for terrestrial small mammal surveying. The Modified Littlewood tunnel produced a higher number of shrew detections compared to the plastic tunnel. The Mostela tunnel produced the highest quality data by facilitating species identification and reducing non-target triggers. Passive acoustic monitoring and camera trapping detected different species, but there was no difference in species richness between the methods. Voles were better detected by camera trapping than passive acoustic monitoring. We consider the success of current machine learning classification for processing our acoustic data, and suggest that future developments in machine learning classification will likely speed up processing of both visual and acoustic data, and perhaps allow more detailed and accurate identification to species and individual level.

## Introduction

Terrestrial small mammals form a large and ecologically important group which, along with many other organisms, are under threat from anthropogenic stressors such as agriculture and land-use change (Byrom *et al*., 2015; Droghini *et al*., 2022; IUCN, 2024). To assess both the impact of stressors and the efficacy of conservation interventions on terrestrial small mammals, we must develop methods suited to long-term, large-scale monitoring of these often-cryptic species.

Traditional wildlife monitoring techniques such as trapping, handling and marking, tissue sampling, and performing manipulation experiments, are considered invasive as they can cause stress and may impact individual growth, reproductive success and survival. Mortensen & Rosell (2020) found that capture and handling of beavers (*Castor fiber*) reduced yearly reproduction and number of kits, and reduced body mass of dominant individuals. A review found that 70% of the 42 mammal species analysed displayed altered behaviour and activity levels after capture and GPS tagging (Stiegler *et al*., 2024).

Invasive methods can also incur several biases – for example, many mammalian species show biased capture rates which vary with age, sex, personality, dominance status and lactation (Buskirk and Lindstedt, 1989; Kay *et al*., 2000; Powell and Proulx, 2003; Way, 2012; Perkins *et al*., 2021; Johnstone, Price and Garvey, 2024). This makes it difficult to gain data representative of a population, and can also alter population structures when kill-trapping is involved. Changes in population sex and age distributions and slight genetic shifts could have negative impacts on the survival of threatened species, which may already have diminished populations and gene pools due to the various aforementioned anthropogenic stressors.

Terrestrial small mammals are traditionally monitored using pitfall traps, or live traps such as Longworth or Sherman traps. Live trapping allows individuals to be physically examined, marked, and samples of fur, toenail clippings, and faeces can be taken. It is therefore possible to obtain data on age, sex, body mass and genetics (Ostfeld *et al*., 2018; Romairone *et al*., 2018), disease and parasite load (Ostfeld *et al*., 2018; Wu *et al*., 2018), microbiome (Kohl, Luong and Dearing, 2015; Stothart, Palme and Newman, 2019; Čížková *et al*., 2021), diet and trophic level (Galetti *et al*., 2016; Ribeiro *et al*., 2019; Balčiauskas, Balčiauskienė, *et al*., 2021), and environmental pollutants (Fritsch *et al*., 2022; Thrift *et al*., 2022). Live trapping is particularly useful as it allows individuals to be marked for recapture studies by fur snipping or ear punching, allowing individual identification and therefore accurate population estimates such as density and abundance can be obtained. Individuals can also be reidentified by using tags such as GPS collars, passive infrared transponder (PIT) tags, or radio transmitters, but micro-versions must be used for mammals with small bodyweights (Lennox *et al*., 2016). Several studies have demonstrated that such micro-attachments can be affixed to and used on terrestrial small mammals (Rychlik, Ruczyński and Borowski, 2010; Collins and Kays, 2014; McMahon *et al*., 2017; Dutt, Veals and Koprowski, 2020; Beatham *et al*., 2021; Hummell, Li and Mullinax, 2022; Wallace *et al*., 2022; Robinson *et al*., 2024; van der Putten *et al*., 2025), although limited studies have tested whether these tags affect long-term stress, behaviour, survival or reproduction. Some research has suggested that there may be negative impacts on activity levels (Hamley and Falls, 1975), social behaviour (Berteaux, Duhamel and Bergeron, 1994), mortality (Wolton and Trowbridge, 1985), and reproduction (Moorhouse and Macdonald, 2005), however other studies find no effect (Berteaux *et al*., 1996; Bolduc *et al*., 2022).

Animals can also be reidentified by mark-release-recapture programs using live trapping. However, these programs can be fieldwork intense, often requiring hundreds of trap nights, sometimes with multiple trap checks a day (Castañeda *et al*., 2018; Harkins, Keinath and Ben-David, 2019; Weldy *et al*., 2019; Freeman *et al*., 2022). In addition, live-trapping can involve animals being left in traps for hours before being examined and released, which is traumatic and can cause death (Romairone *et al*., 2018; Littlewood *et al*., 2021). Given the need to reduce handling time and trauma to the animal, traps can only be monitored by trained persons. The method also incurs some biases towards sexes, individuals, or species (Hammond and Anthony, 2006) which are predisposed to being re-trapped, for example ‘trap-happy’ individuals who learn they will be caught but released, and therefore return to eat the bait (Torre *et al*., 2016; Newson and Pearce, 2022).

Passive, minimally-invasive, or non-invasive technologies, collect data with little or no disturbance to the usual behaviours, ecology and physiology of organisms and ecosystems. Given the reduced impact that non-invasive methods have on individuals and environments, monitoring can reasonably be longer-term and larger-scale. This is particularly useful for use in sensitive habitats (McCleery *et al*., 2022) and in monitoring cryptic, rare and threatened species (Caravaggi *et al*., 2017). Non-invasive methods do not usually require constant monitoring as live traps do, and so can be left untended for longer periods. Reducing field time means lower costs, and is particularly helpful in surveying remote, inaccessible or dangerous environments, such as remote islands (Dueser, Porter and Moncrief, 2025), cliff faces (Newson *et al*., 2023), regions of extreme climatic conditions or difficult terrain (McDonald *et al*., 2015), arboreal (Moore *et al*., 2021), subnivean (Soininen *et al*., 2015) and subterranean habitats (Lisse and Pinot, 2024). Technological advancements allowing data to be uploaded from field equipment via satellite, and self-powered equipment (Corva *et al*., 2022; Whytock *et al*., 2023; Pestell *et al*., 2025) will further the reduce field time and ecosystem disturbance from passive techniques. Reducing field time could make such methods more efficient, providing that these methods do not require extra time at other stages; this will be discussed later.

Non-invasive wildlife monitoring methods include thermal imaging (Jerem *et al*., 2018; Mapes *et al*., 2020; Witt *et al*., 2020; Underwood, Derhè and Jacups, 2022), camera trapping (McCallum, 2013; Trolliet *et al*., 2014; Burton *et al*., 2015; Meek, Ballard and Fleming, 2015; Caravaggi *et al*., 2017; Frey *et al*., 2017; Jumeau, Petrod and Handrich, 2017; Steenweg *et al*., 2017; Delisle *et al*., 2021; Gilbert *et al*., 2021; Hereward *et al*., 2021; Moore *et al*., 2021; Preti, Verheggen and Angeli, 2021; Barroso and Palencia, 2025), acoustic monitoring (Sugai *et al*., 2019; Pérez-Granados and Traba, 2021; Fleishman *et al*., 2023; Hoefer *et al*., 2023; Ross *et al*., 2023; Sharma, Sato and Gautam, 2023), detection dogs (Leigh and Dominick, 2015; Cristescu, Miller and Frère, 2020; Karp, 2020), and analysis of eDNA (Thomsen and Willerslev, 2015; Leempoel, Hebert and Hadly, 2020; Lyet *et al*., 2021; Mena *et al*., 2021; Schwentner *et al*., 2021), faeces (Oja *et al*., 2017; Mengüllüoğlu *et al*., 2019; Davey *et al*., 2023), and roadkill (Balčiauskas, Stratford, *et al*., 2021; Thomas *et al*., 2022). Many non-invasive methods have been applied to studying small mammals, including analysis of faeces (Torre *et al*., 2013; O’Meara *et al*., 2014; Vigués *et al*., 2021) and owl pellets (Heisler, Somers and Poulin, 2016; Stutz *et al*., 2020; Schoenefuss *et al*., 2024), and use of detection dogs (Thomas *et al*., 2020), hair tubes (Pocock and Jennings, 2006; Mortelliti and Boitani, 2008; Mortelliti *et al*., 2010; Chiron *et al*., 2018; Dürger *et al*., 2024), footprint tunnels (Palma and Gurgel-Gonçalves, 2007; Atkins *et al*., 2018; Harrow, Horncastle and Chambers, 2018), camera trapping (Soininen *et al*., 2015; Hobbs and Brehme, 2017; Amber, Lipps and Peterman, 2020; Littlewood *et al*., 2021) and recently, bioacoustics (Newson, Middleton and Pearce, 2020; Newson and Pearce, 2022).

Some of these methods allow researchers to obtain genetic, behavioural and presence data, but may not be best suited to all studies. Analysis of faeces and hair allows unintrusive collection of genetic data, but gene sequencing techniques can be expensive (Lancaster *et al*., 2023; Theissinger *et al*., 2023) and therefore are best utilised when genetic data is essential, for example in genetics studies or in distinguishing morphologically similar species. Footprints obtained using tracking tunnels can have variable detection rates (Wiewel, Clark and Sovada, 2007), and can be difficult to classify to species without expertise (Glennon, Porter and Demers, 2002; Wiewel, Clark and Sovada, 2007; Brehme *et al*., 2019; Yiu, Etherington and Russell, 2025), although computerised methods can help with this (Palma and Gurgel-Gonçalves, 2007; Russell *et al*., 2009; Kistner *et al*., 2024; Tucker *et al*., 2024). Analysis of owl pellets is naturally biased by the owl’s choice of prey species (Torre, Arrizabalaga and Flaquer, 2004; Comay and Dayan, 2018), and can also require expertise where many closely related species coexist (Charley, Gray and Baker, 2025). Development of a broader selection of non-invasive methods would allow optimal choice of a method based on study aims and budget.

Camera trapping is a non-invasive technique usually involving deployment of motion-triggered cameras, which record videos or still photographs of animals in the area. Camera trapping has proved extremely useful and adaptable, being used on birds (O’Brien and Kinnaird, 2008; Randler and Kalb, 2018; Fontúrbel *et al*., 2020), herpetofauna (Ryberg *et al*., 2021; Antunes *et al*., 2022; Brown, Hannon and Maerz, 2023; Olson, Laughlin and Martin, 2023; Acker-Cooper, Roxburgh and Tarrant, 2025), and mammals (McCallum, 2013; Rovero *et al*., 2014; Chen *et al*., 2022; Cordier *et al*., 2022; Hsing *et al*., 2022) across habitats ranging from open grassland to floodplain forests (Swanson *et al*., 2015; Mere Roncal *et al*., 2019). Camera trapping has been shown to be as or more effective than other methods including live-trapping (Wearn and Glover-Kapfer, 2019). While camera trapping for terrestrial small mammals does require some adaptations, many studies have addressed these issues and developed successful adjustments. A primary adaptation needed when moving from recording large (*e.g.* deer, big cats etc.) to small mammals is the ability to focus on a small target at close range. This can be achieved by placing cameras within boxes or facing the ground from a post, and/or fitting close-focus lenses over the cameras’ in-built lenses to a) reduce the focal distance relative to the subject species and b) narrow the field of view (Rendall *et al*., 2014; Croose and Carter, 2019; Littlewood *et al*., 2021).

Passive acoustic monitoring (PAM) may also provide a useful alternative to some invasive methods. PAM is a non-invasive form of bioacoustics, which collects sound data from the environment, including infrasonic sounds below, sonic or audible sounds within, and ultrasonic sounds above, the range of human hearing (20Hz-20kHz) (Payne, Langbauer and Thomas, 1986; Schnitzler and Kalko, 2001; Mellinger and Clark, 2003; Jones and Holderied, 2007; Britzke, Gillam and Murray, 2013; Gibb *et al*., 2019; Sugai *et al*., 2019). Bioacoustics are currently being used to monitor species presence and behaviours in marine animals (King, Connor and Montgomery, 2022; Cominelli *et al*., 2024), bats (López-Baucells *et al*., 2021; Froidevaux *et al*., 2023; Mancini *et al*., 2024, 2024) and, more recently, terrestrial mammals (Barber-Meyer *et al*., 2020; Penar, Magiera and Klocek, 2020; Pérez-Granados and Schuchmann, 2021). Terrestrial small mammals are known to produce audible and ultrasonic vocalisations which are used in communication and echo-orientation (Dent, 2018; Chai *et al*., 2020; Barker *et al*., 2021; Warren *et al*., 2025). While mostly not well documented, evidence suggests that many small mammal species have various structurally complex functional call types (Sales, 2010; Newson, Middleton and Pearce, 2020). Acoustics may be helpful in detecting cryptic species (Ancillotto *et al*., 2017; Diggins *et al*., 2020; Volodin *et al*., 2022), and distinguishing morphologically-similar species (Ancillotto *et al*., 2017); frequency analysis of the echolocating calls of pipistrelle bats in Britain revealed that two distinct species are present rather than one – common pipistrelle (*Pipistrellus pipistrellus*) and soprano pipistrelle (*P. pygmaeus*) (Jones and Van Parijs, 1993; Barratt *et al*., 1997; Jones and Barlow, 1999). Several species’ vocalisations have been studied and used in monitoring (Barščevska, Juškaitis and Adamík, 2024; McEwen *et al*., 2024; Vilalta, Marina Rodríguez, 2024) however this is not yet widely used across species, and there is so far a lack of understanding of how bioacoustics compare to other monitoring methods (Newson and Pearce, 2022).

Given that both camera trapping and PAM generate complementary records of data, the techniques should therefore complement each other in revealing aspects of animal behaviour and ecology. PAM and camera trapping have so far only been directly compared in studies on chimpanzees (*Pan troglodytes*) (Crunchant *et al*., 2020), sika deer (*Cervus nippon*) and Japanese macaques (*Macaca fuscata*) (Enari *et al*., 2019); these studies found that PAM had a much larger detection distance (between 17 and 1,738 times larger) than camera trapping. However, this may not hold true for small mammals, whose lower amplitude, higher frequency calls are likely to travel much shorter distances. These comparative studies have mainly focussed on detection distance but future studies should compare cost, processing time, and accuracy, as well as performing these studies on a wider range of taxa including small terrestrial mammals. PAM, camera trapping and GPS collars were combined in a study on predator-prey interactions which looked at the response of leopards to monkey alarm calls (Isbell and Bidner, 2016), demonstrating that there is significant potential for use of PAM and camera trapping in studying animal behaviour and communication. Mine *et al*. (Mine *et al*., 2024) combined visual and auditory surveys to study multi-modal communication in chimpanzees. PAM and camera trapping have also been paired to look at the effect of tourist noise on cormorant disturbance behaviours at nest sites (Buxton *et al*., 2017). Behavioural studies can provide information on species’ habitat requirement, dispersal and reproductive behaviour, which can help in deciding which interventions are needed and how conservation projects can maximise positive impacts (Caravaggi *et al*., 2017).

Camera trapping therefore has yet to be compared to continuous acoustic monitoring for small mammals. Studies attempting this should compare number of detections, both the percentage of camera trap detections during which the animals also vocalise, and the percentage of visits detected by PAM that are also detected by camera traps.

Here, we compare the performance of three small mammal camera trapping tunnel designs to investigate whether one tunnel is better suited to (British) terrestrial small mammal monitoring than the others. To address this, we explored whether the tunnel designs differed in small mammal detection frequency and in the latency to first detection of species group. We also explored whether quality parameters of the visual data obtained, such as rate of species identification, or number of non-target triggers, differed between the tunnel designs. We also assess the potential of passive acoustic monitoring to be used for small mammals by comparing it to camera trapping, a more established and understood small mammal monitoring technique. To do this, we explore whether acoustic recorders and cameras are able to detect the same species when they are deployed together. Finally, we aim to assess whether the use of passive acoustic monitoring and camera trapping together are able to generate meaningful information about social behaviour, by asking whether small mammals vocalise more frequently when multiple individuals are present. In this study, ‘terrestrial small mammal’ refers to non-volant mammals under 5kg.

## Methods

The fieldwork described in this study was reviewed and approved by the University of Bristol Animal Welfare Ethical Review Board (University Investigation Number UIN-25-008).

### Study Areas

Surveys were conducted at four sites in the United Kingdom: three sites near to Bristol in south-west England, and one site in south Wales (Figure 1). These sites included a variety of non-urban habitats, so that the results were relevant to a range of habitat types. Fenswood Farm (located in North Somerset, surveyed May – June 2024) is an arable farm (wheat, oats, mustard) with some cattle grazing managed by the University of Bristol. Lower Woods (located in South Gloucestershire, surveyed July – August 2024) is a nature reserve and Site of Special Scientific Interest managed by Gloucestershire Wildlife Trust. Great Avon Wood (located in North East Somerset, surveyed August – September 2024) is a forest regeneration project on land that was farmed until 2023, and is managed by Avon Needs Trees. A smallholding farm in the Black Mountains, Bannau Brycheiniog National Park, South Wales, was surveyed in June 2024. Fuller descriptions of the sites, including previous sampling history, land use, vegetation, and environmental data during the study are given in Appendix 1.

**Figure 1.**
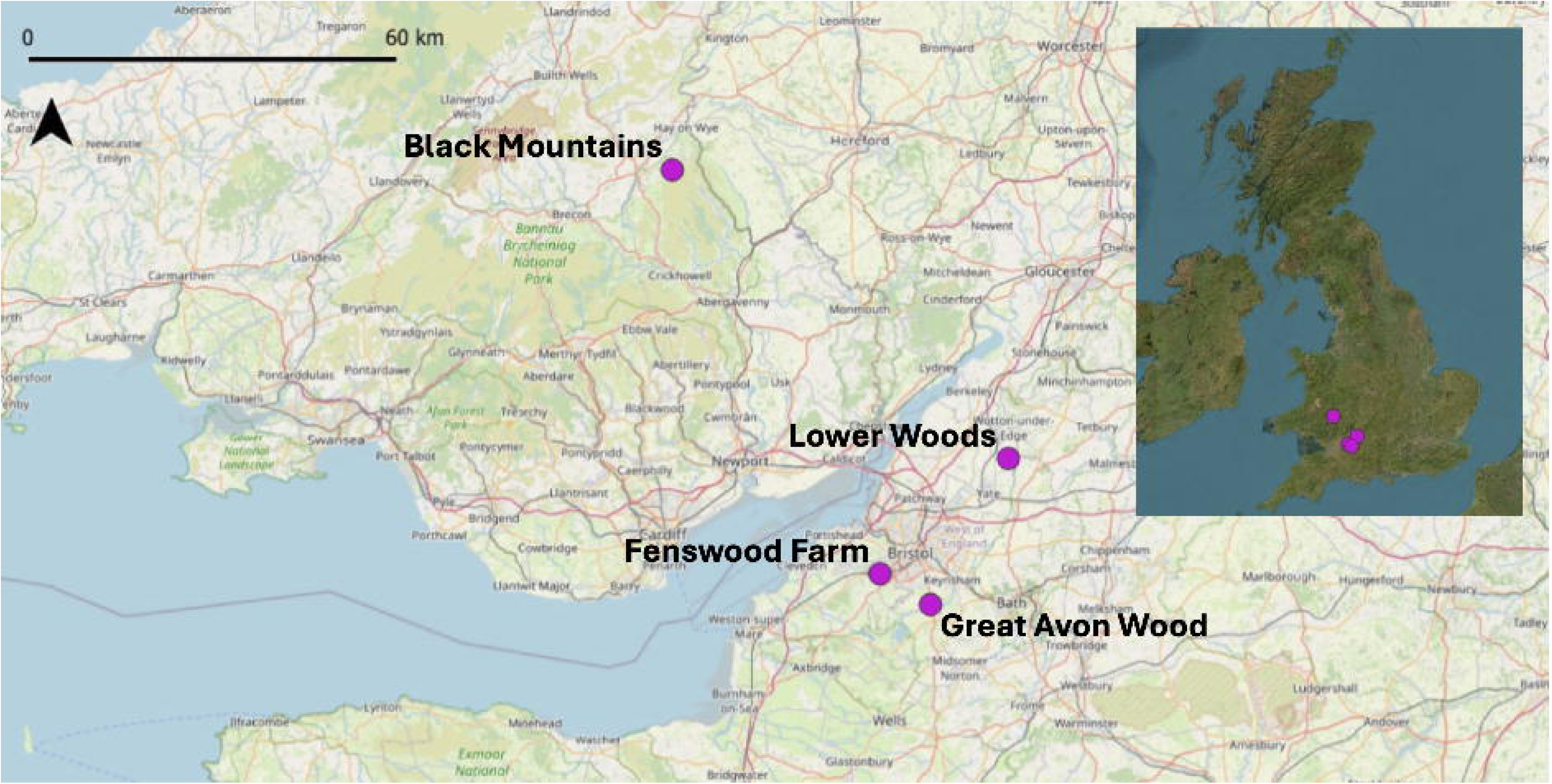
Locations of four sites chosen for small mammal monitoring in Britain, including one site in South Wales, and three sites in South West England. Map produced using QGIS (QGIS Development Team, 2025).

## Materials and Equipment

### Tunnel Designs

#### We considered three tunnel designs

Modified Littlewood: we adapted the design described by Littlewood *et al*. (Littlewood *et al*., 2021) (Figure 2), constructing the tunnels using recycled softwood decking planks. Unlike the original design, which included a Perspex roof, our tunnels had wooden roofs, which may offer greater shelter to visiting species. The Modified Littlewood tunnels had been in use for one year before this study, so despite being cleaned between uses, they may have retained smells of previous small mammal visitations, and any smells of wood treatment were likely diminished. Modified Littlewood tunnels were fitted with a camera trap at the end of the tunnel, opposite the entrance (Figure 2A).

**Figure 2.**
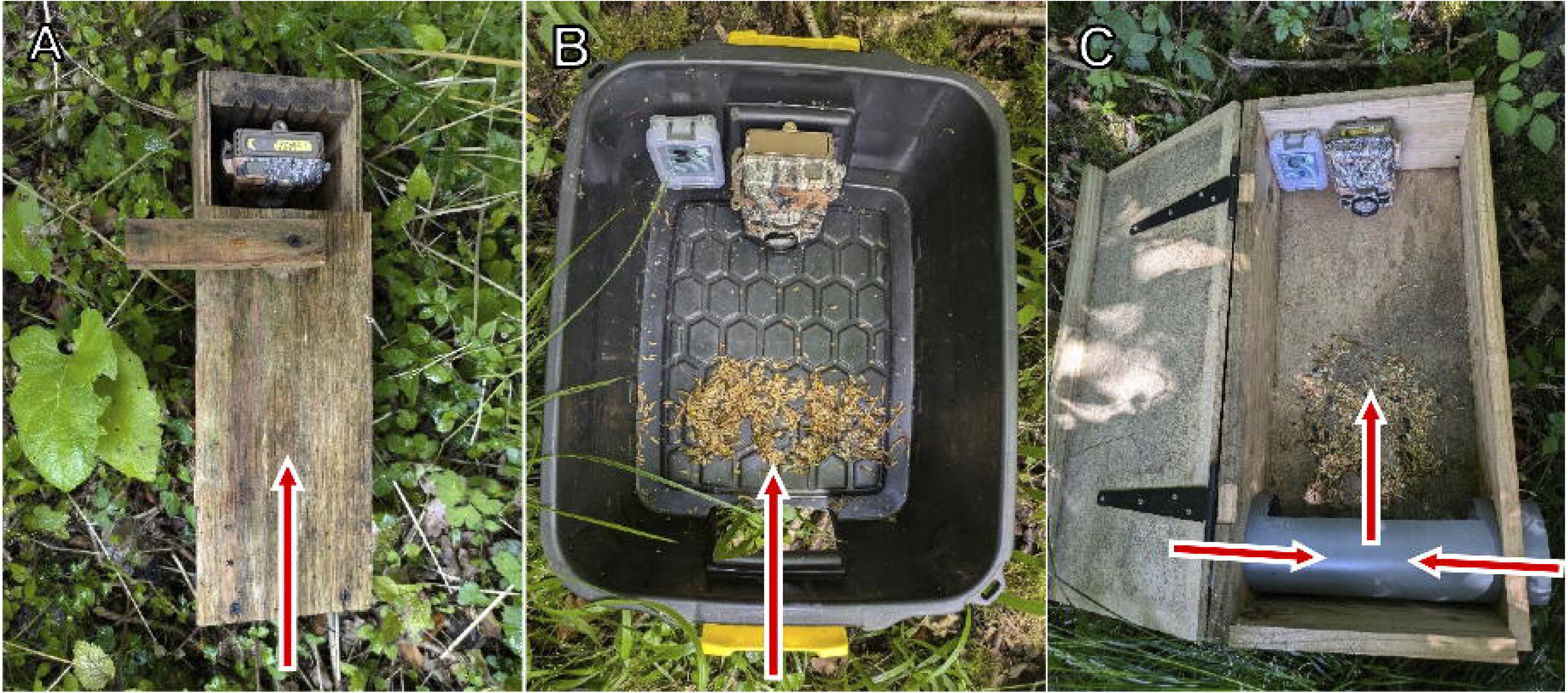
Three designs of terrestrial small mammal monitoring tunnels (without lids). **A**: Modified Littlewood tunnel containing a trail camera; **B**) plastic tunnel containing a trail camera and an acoustic monitor; **C**) Mostela tunnel containing a trail camera and an acoustic monitor. Red and white arrows indicate entrances/exits for small mammals.

Plastic: to explore a trap that could be constructed quickly with simple, inexpensive off-the-shelf products, we designed these tunnels using domestic plastic storage boxes (95% recycled plastic 24L Strata storage boxes, Homebase, Plymouth). Modifying the storage boxes took approximately ten minutes per box using a retractable utility knife. Plastic tunnels housed a trail camera and an acoustic recorder (Figure 2B).

Mostela: these tunnels were based on a design created for camera trapping mustelids in the Netherlands (Croose and Carter, 2019). Tunnels were newly built for this study using marine ply for the floor and roof, pine plank sides, and a PVC pipe (Figure 13). We used 10 cm diameter PVC pipe as this was found to produce a higher detection probability of weasels and stoats compared to 8 cm diameter pipe (Mos and Hofmeester, 2020). Mostela tunnels housed a trail camera and an acoustic recorder (Figure 2C).

We used Spec Ops Advantage cameras and Spec Ops Edge cameras (Browning Trail Cameras, Birmingham, USA) both of which use no-glow infrared flash, and were configured with identical settings (Appendix 2). The Spec Ops Advantage cameras had a fast trigger speed of 0.4 seconds, and Spec Ops Edge cameras had a fast trigger speed of 0.2 seconds, so cameras were randomly allocated to tunnels each deployment. Initially five Browning Spec Ops Edge and three Browning Spec Ops Advantage were used, however after two weeks one Spec Ops Advantage failed, and was replaced by a Spec Ops Edge. 37 mm diameter +4 close focus camera lenses (Fotover macro filter kit) were attached in front of the camera lenses using adhesive putty (Blu Tack®). +1, +2, +4, and +10 macro lenses were tested, with +4 producing the highest resolution images at the correct focal distance. Cameras and acoustic recorders were randomly allocated to tunnels before each deployment.

We chose to record videos, rather than capture still images. Recording videos is less efficient than capturing still images, producing similar detection rates (Glen *et al*., 2013), but requiring more storage and processing time (Glen *et al*., 2013; Smaal and van Manen, 2022). Several studies have found no difference in ability to perform species identification between videos and still images (Glen *et al*., 2013; Taylor *et al*., 2013; Smaal and van Manen, 2022). However, Green *et al*. (Green *et al*., 2023) found that while identification ability did not differ for expert identifiers, citizen scientist classifications were more accurate for videos than photos. We were also interested in exploring whether behaviours could be captured accurately with the tunnels, which requires video footage.

Acoustic recorders were fitted into the Plastic and Mostela tunnels (the Modified Littlewood tunnels did not have sufficient space). We used AudioMoth acoustic recorders version 1.2.0 (https://www.openacousticdevices.info/audiomoth), which were all configured with identical settings with AudioMoth-Firmware-Basic version 1.10.0 (Table A2). Full detail of configuration and the containers used to house the recorders are given in Appendix 3.

## Fieldwork

### Site Selection

A map of each site was divided into a grid of 200 × 200 m. This distance was chosen to reduce the number of individuals travelling between deployments, and was decided based on information regarding home ranges of rodents and shrews (Borowski, 2003; Godsall, Coulson and Malo, 2014; Frafjord, 2016; Boratyński, Szyrmer and Koteja, 2020). Mustelids were not a primary focus of the study due to low detection rates and the time constraints of this study and were therefore not considered in this selection process. Unless deemed unsuitable for surveying, each square on the site was given a number and squares were randomly selected before each deployment. Squares were deemed unsuitable if they contained an inhabited building or area such as a car park, or if cameras would be at risk of theft due to popular use by the public. For each deployment period, three squares were chosen at random and three triplets containing one of each tunnel design were deployed in the centre of each square.

### Tunnel Deployment

Deployments were performed in triplets, with each triplet containing one of each tunnel type. Within a triplet, all tunnels were deployed containing a randomly selected camera, and the plastic tunnel and Mostela tunnel also contained a randomly selected AudioMoth. Within a triplet, tunnels were deployed approximately 20 m apart, either in a line or in a triangle formation (Figure 3). A triangle was deemed best as all tunnels would then be equidistant, removing bias from the central tunnel receiving more visitors. However, when hedges, rivers or other linear features were present, tunnels were placed in a line, with entrances all facing the same direction. This resulted in 17 linear deployments and 8 triangular deployments.

**Figure 3.**
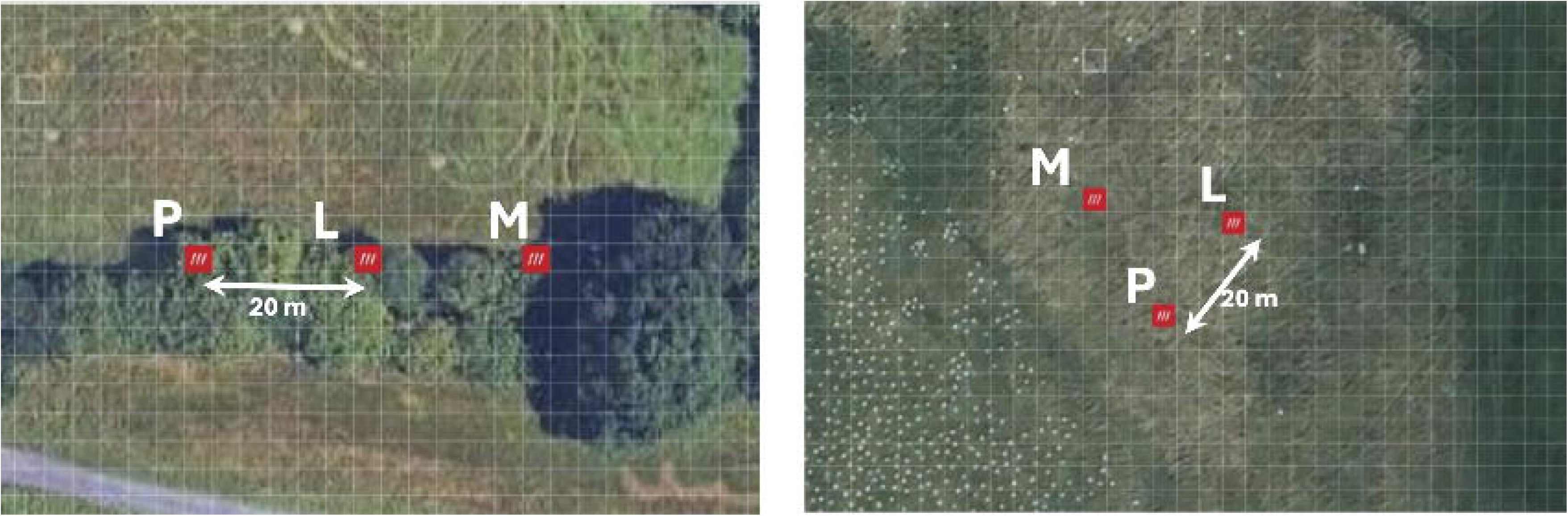
Terrestrial small mammal camera trapping tunnels were deployed in a random order and were either deployed in a triangle formation or a line formation if linear landscape features were present. Tunnels within these triplets were deployed approximately 20 m apart, as shown by white arrows. Tunnel designs were the Modified Littlewood tunnel (L), the Mostela tunnel (M), and the plastic tunnel (P). Screenshots taken from What3Words (https://what3words.com/).

Doing so exposed each tunnel to a similar microhabitat, as microhabitat is shown to influence small mammal communities (Mengak and Guynn, 2003; Churchfield and Rychlik, 2006; Fay, Banks, B. and Korpimäki, 2006; Keckel, Ansorge and Stefen, 2014; Sanches *et al*., 2023; Ancillotto *et al*., 2025). Within these lines or triangle formations, tunnels were deployed in a random order and position with respect to the other tunnel designs. Tunnels were a minimum of 170 m away from tunnels in separate triplets.

AudioMoths were secured to inside of the tunnel wall using Velcro. Cameras were deployed upside down to obtain the desired field of view. All tunnels were baited with dried mealworms and mixed seed (wheat, cut maize, bruised oats, red millet, black sunflower, rolled barley, veg oil) (Castle Bird Feeds, Wild Bird food). Bait was used to attract the focal animal into the optimal range for detection and production of a high-quality video for identification (Tennant *et al*., 2020; Gracanin, Minchinton and Mikac, 2022). Little research has focused on the effect of attractants on small mammal detection rates (Stickel, 1948; Holinda, Burgar and Burton, 2020) other than small carnivores (Mills *et al*., 2019; Holinda, Burgar and Burton, 2020; Randler *et al*., 2020; Davis, Saloojee and Venter, 2025); however, Randler *et al*. (Randler *et al*., 2020) show that use of attractants can facilitate species identification by keeping the animal in the detection area for longer.

Each tunnel was left undisturbed for 7 nights, or approximately 168 hours. At this point, recording was ended, all data were removed from the SD cards, and any remaining bait was removed from site. Tunnels were cleaned after each deployment using a scraper to remove animal scat and seed shells. For each new deployment, fresh bait and batteries were used and SD cards were reformatted to reduce transfer errors.

Two field days were required each week, one to deploy the tunnels and one to collect the SD cards and batteries. Time required to deploy or collect tunnels varied depending on the site and distance between triplets, but took approximately four hours, excluding travel time to the site. Therefore around 8 hours of on-site fieldwork were required per week to deploy three triplets, and 72 hours total were spent over the 9 weeks of fieldwork.

During the study, equipment failure affected one AudioMoth and three camera traps. Camera trap data was incomplete due to early battery failure on one deployment. Disturbance by humans and other animals caused three additional tunnels’ video data to be omitted.

## Data processing

### Visual data

Complete triplets of video data were subset and manually processed to identify species, time of recording, and number of individuals. Incomplete triplets, where one or more of the three tunnel designs did not have a complete set of video data, were not included in subsequent analyses. Incomplete triplets resulted from animal interference, cameras failing to record for the entire duration, or from some or all videos being lost or corrupted during data transfer from the SD card. If no animal was present in a video, it was marked as a non-target trigger, and the suspected reason for the trigger was recorded – reasons included triggers by large mammals, birds or invertebrates, or weather causing moving vegetation. If no small mammal was visible during the video but it was suspected that a small mammal was off-frame due to noise of eating, the video was still marked as a non-target trigger, as we were comparing the visual data from camera trapping to bioacoustics. If an individual spent over two seconds off-frame, this was also recorded; an individual was considered off-frame if the view of the animal’s body was insufficient to allow species identification or classification of behaviour.

Animals were identified to species level when possible with full confidence, or if not, genus. We did not attempt to distinguish wood mice (*Apodemus sylvaticus*) and yellow-necked mice (*A. flavicollis*), as they can only be reliably distinguished by inspection of the yellow band on their throat. The majority of videos were in black and white, and the throat was not usually seen. Given that both species are present in the UK, they were automatically classified as *Apodemus* spp. Books and online resources were used to assist animal identification, including *Tracks and Signs of the Animals and Birds of Britain and Europe* (Olsen and Epstein, 2013) and the *Mammal Mapper* app (Mammal Society: https://mammal.org.uk/mammal-mapper).

### Acoustic data

A subset of the data was made containing deployments with complete audio and video data. Datasets were considered to be incomplete when one or more acoustic or video file was lost or corrupt within that AudioMoth’s battery lifespan, which averaged 57 hours. Datasets including lost or corrupt files after the AudioMoth had stopped recording were still considered complete, as only data within AudioMoth battery lifespan was used in this analysis. Deployments where no videos of small mammals had been recorded within the AudioMoth battery lifespan were also excluded from this subset.

Audio files were then subsetted according to their timestamps. All acoustic recordings occurring within two minutes of any video recording were copied to another folder for further investigation. A two-minute window was used to account for potential discrepancies in the internal clocks of trail cameras and AudioMoths, ensuring that temporally matched events were not missed due to mismatched time keeping between devices. Subsetting of acoustic files was performed for all temporally matched videos, including non-target triggers, to account for the fact that non-target triggers can be caused by the camera triggering too slowly to catch an animal just leaving. Analysis of acoustic recordings which did not occur at the same time as a video would have provided useful data on camera trap false negatives, however this study was constrained by the limited free data allowance of the acoustic processing software used.

This subset of files which had been recorded within two minutes of a video were then processed using the British Trust for Ornithology’s Acoustic Pipeline version 5.611 (British Trust for Ornithology: https://www.bto.org/data/tools-products/acoustic-pipeline). Small mammal classifications of below 0.5 confidence level were then manually checked to validate the Acoustic Pipeline’s output. Manual verification involved visualisation of spectrograms in Kaleidoscope lite software version 5.6.8 (Wildlife Acoustics, Inc., Maynard, MA, USA), and reference to Middleton (Middleton, 2020), Middleton *et al*. (Middleton, Newson and Pearce, 2023), and Newson *et al*. (Newson, Middleton and Pearce, 2020).

At the time of use, the *BTO Acoustic Pipeline*’s small mammal coverage did not include mustelids, lagomorphs, or squirrels, therefore for the acoustic data we only considered voles, rats, mice, dormice, and shrews.

## Data analysis

All data and associated *R* code are freely available at *Zenodo* (Stiff, Wakefield and Rands, 2026).

### Visual data

All analysis was performed on species groupings of mice, of voles, and of shrews, rather than individual species, as in the video data, field voles and all shrew species had low counts, and mice were rarely identified to species level. All statistical analyses were performed in *R* 4.1.2 (R Development Core Team, 2025). Packages used were *glmmTMB* (Brooks *et al*., 2017), *lme4* (Bates *et al*., 2015), *lmerTest* (Kuznetsova, Brockhof and Christensen, 2017), *ggplot2* (Wickham, 2009), *emmeans* (Lenth *et al*., 2025), *moments* (Komsta and Novomestky, 2022), *dplyr* (Wickham *et al*., 2023), *performance* (Lüdecke *et al*., 2021), *DHARMa* (Hartig, 2025), *FSA* (Ogle *et al*., 2025), *MASS* (Venables and Ripley, 2002), *multcomp* (Hothorn, Bretz and Westfall, 2008), and *purrr* (Wickham and Henry, 2025).

Count data of each taxonomic group per tunnel deployment were extracted from the main database. Camera trap count data were fitted with a generalised linear mixed model to test whether tunnel design influenced data quality or the number of visitations of voles, mice or shrews. Data were not normally distributed, and were strongly left-skewed due to a large number of zeros. Log transformation did not result in a normal distribution, so we used non-parametric tests. We compared the AIC, residual values, and dispersion of Poisson and negative binomial distributions, and found that the data best fit a negative binomial distribution for all small mammal counts and for all data quality measures – non-target triggers, unidentifiable small mammals, and videos where an individual was off-frame. We also checked for over-dispersion and zero inflation, which were not observed.

Given the form of the data, we modelled all count data using negative binomial generalised linear mixed models, with tunnel design as a fixed effect and site and nested deployment as random effects. Pairwise comparisons of tunnel designs were conducted using estimated marginal means (EMMs) with Tukey adjustment for multiple comparisons.

Latency to detection was calculated as the mean number of hours from camera deployment until the first detection of each species group in each tunnel type, in each deployment. For mice and voles, latency to detection was compared between tunnel designs using linear mixed models with the identity of the triplet deployment as a random effect, using *lmerTest*. Mouse data were initially non-normally distributed, so various transformations were compared, with log transformation found to best normalise the distribution of the residual values. Vole data were already normally distributed and therefore no transformation was performed. Pairwise comparisons were conducted using estimated marginal means with Tukey adjustment for multiple comparisons.

For analysis of latency to detection of shrews, only the Modified Littlewood (*n* = 7) and Mostela (*n* = 5) tunnels were included, as the plastic tunnel only detected shrews on two deployments. The shrew data were not normally distributed and transformations were unsuccessful. Given this and the small sample size, a Wilcoxon signed rank test was used to compare latency to first detection of shrews for the Modified Littlewood and the Mostela tunnels.

### Acoustic data

The presence or absence of each species group was recorded for each triplet deployment, for both camera trapping and acoustic monitoring, during deployment time when both camera trap and AudioMoth were active. To test whether PAM and camera trapping produced similar results of species groups detected, an exact McNemar test was performed on paired binary data of presence and absence of each species group detected by PAM and camera trapping, each deployment/each tunnel type deployment. Species groups included were mice, rats, shrews, voles, and dormice. The McNemar test is based on the number of discordant pairs – cases where the outcome differs between the two methods for the same tunnel deployment. According to Pembury Smith & Ruxton (Pembury Smith and Ruxton, 2020), at least four discordant pairs are required in a McNemar test for a meaningful *p* value, and that large effect sizes are needed to produce significant results with fewer than ten discordant pairs, due to low statistical power. Therefore, we also performed an exact McNemar test on individual species groups with more than four discordant pairs, which was only the vole data. Mice, rats, dormice, and shrews each had fewer than four discordant pairs and therefore they were not tested separately.

### Behavioural data

Initially, we planned to assess whether vocalising was associated with any specific behaviours seen on video (such as foraging or grooming), however due to the small number of small mammal classifications by the pipeline, this was not possible. Instead, we analysed whether small mammal calling was associated with social behaviour, using two tests.

Of complete audio-video deployments, paired data were extracted on whether each species had been detected by PAM or camera trapping, and whether multiple individuals were seen together in any video within that deployment (as noting that multiple individuals are present in the videos is the only means by which we can confidently state that social behaviour might be possible). To account for interspecific and intraspecific social communication, videos were classed as containing multiple individuals even if the individuals were of different species groups. An exact McNemar test was then performed on these data.

As an additional test, each positive small mammal detection by the BTO Acoustic Pipeline was matched with the video file at the corresponding time. For each deployment where a small mammal vocalisation had been detected by the pipeline, data was extracted on whether multiple individuals had been seen together in one of the audio-matched videos. An exact McNemar test was then performed on these paired data. In all McNemar tests, exact tests were used due to low numbers of discordant pairs.

## Results

### General overview of data

#### Visual Data

Overall, the cameras detected thirteen small mammal taxonomic groupings: field vole *Microtus agrestis*, bank vole *Myodes glareolus*, brown rat *Rattus norvegicus*, house mouse *Mus musculus*, *Apodemus* spp. mouse (wood mouse *A. sylvaticus* or yellow-necked mouse *A. flavicollis*), common shrew *Sorex araneus*, pygmy shrew *S. minutus*, water shrew *Neomys fodiens*, grey squirrel *Sciurus carolinensis*, European hare *Lepus europaeus*, European rabbit *Oryctolagus cuniculus*, least weasel *Mustela nivalis*, and European polecat *Mustela putorius* (Figure 4). Also detected were red fox *Vulpes vulpes*, European badger *Meles meles*, unidentifiable species of deer, and several species of bird and invertebrate. Confidence in distinguishing *Apodemus* spp. from house mice was often low, leading to large numbers of unclassified mice.

**Figure 4.**
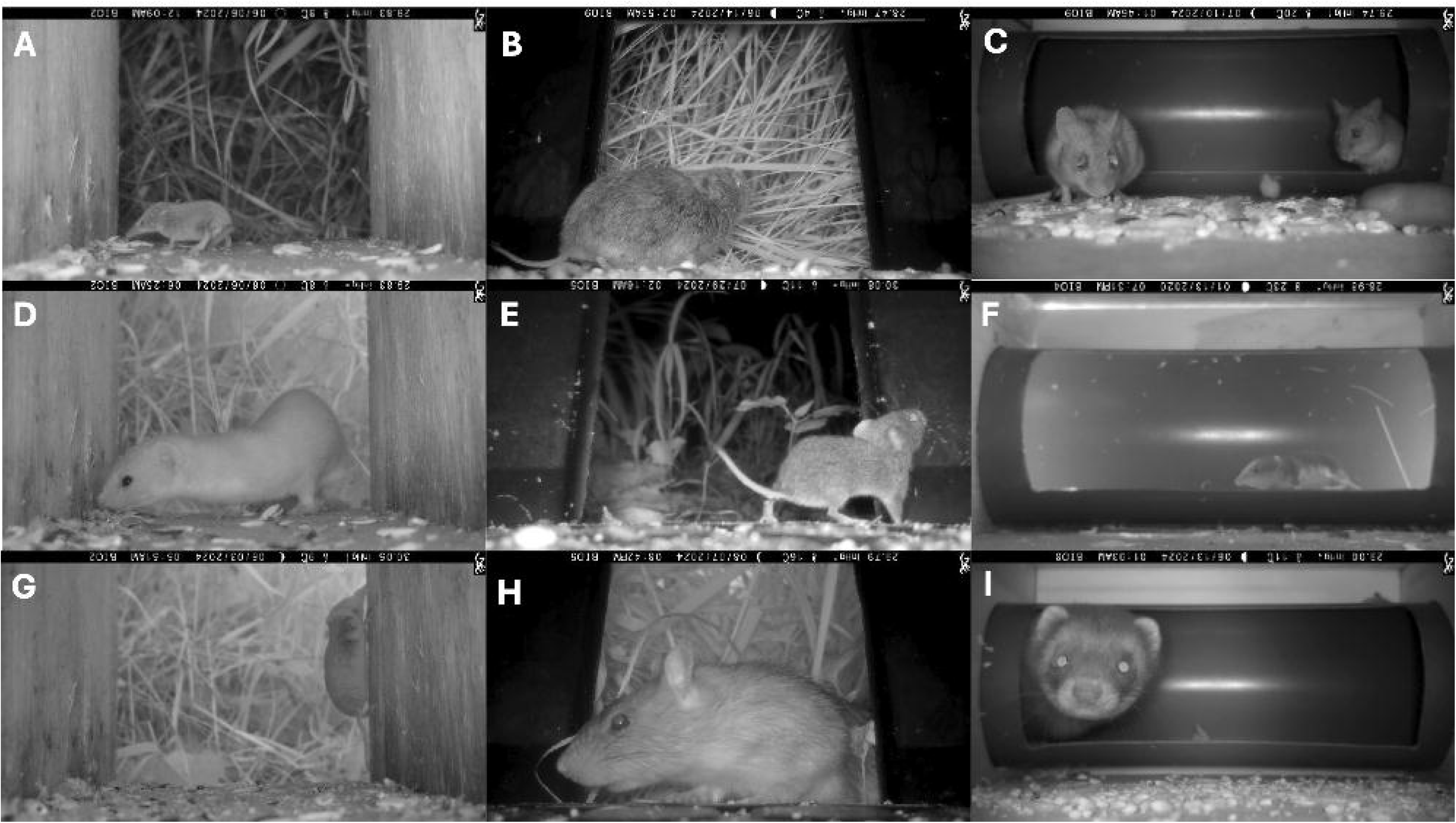
Example images of small mammal species detected through camera trapping using three tunnel types. Images in the left column were taken in the Modified Littlewood tunnel, those in the middle were taken in the plastic tunnel, and those on the right were taken in the Mostela tunnel. Species shown include pygmy shrew (A), field vole (B), *Apodemus* sp. (C), least weasel (D), bank vole (E), water shrew (F), common shrew (G), brown rat (H), polecat (I).

Twenty-six triplets were deployed, which resulted in 16 complete camera triplets (Table 1). The 16 complete video triplets amounted to 11,964 videos collected over 336 trap nights; taking approximately 60 hours to process manually. Of these complete triplet videos, 9,112 (76%) videos contained a small mammal, the rest were non-target triggers caused by moving vegetation, large mammals, birds, invertebrates, or human disturbance. 3.2% of small mammals could not be identified to the level of mouse, shrew, vole, mustelid, squirrel.

**Table 1.** Deployment of camera traps and AudioMoths targeting terrestrial small mammals. One triplet includes the deployment of one Modified Littlewood tunnel (containing a trail camera), one plastic tunnel (containing a trail camera and an AudioMoth), and one Mostela tunnel (containing a trail camera and an AudioMoth). A complete camera trap triplet was obtained when all three cameras within the triplet recorded for the full duration of the deployment, and no files were lost or corrupt. A complete audio-visual pair was obtained when the camera and the AudioMoth from the same tunnel (of any design) produced complete data, with no audio or video files lost or corrupt within the AudioMoth’s lifespan, and with at least one small mammal detections within the Audiomoth battery’s lifespan.

| Study area | Deployment period | Tunnel triplets deployed | Complete camera trap triplets obtained | Complete audio-visual pairs obtained |
| --- | --- | --- | --- | --- |
| Fenswood Farm | May – June 2024 | 6 | 3 | 5 |
| Black Mountains | June 2024 | 3 | 2 | 0 |
| Lower Woods | July – August 2024 | 9 | 5 | 9 |
| Great Avon Wood | August – September 2024 | 8 | 6 | 6 |
| Total |  | 26 | 16 | 20 |

Detection of different small mammal groups varied among sites, with the Black Mountains having highest detection rates for mice and shrews, and Fenswood Farm having the highest detection rates for mustelids and voles (Table 2). Numbers of small mammal detections also varied across tunnel types (Table 3).

**Table 2.** Differences in terrestrial small mammal detections by camera trapping across four sites in Britain. Number of camera deployments varied across sites therefore average detections are per camera are given rather than raw figures. A ‘detection’ refers to a video recording rather than an individual small mammal, therefore calculations do not account for multiple small mammal visits during one video. Small mammal detections includes mice, voles, shrews, mustelids, rats, and squirrels. Figures are calculated from all processed complete tunnel deployments, not only complete triplet deployments.

| **Average detections per camera deployment** | **Fenswood Farm** | **Black Mountains** | **Lower Woods** | **Great Avon Wood** | **Overall average** |
| --- | --- | --- | --- | --- | --- |
| **Overall small mammals** | 83.8 | 231.9 | 221.8 | 176.1 | 178.4 |
| **Mice** | 25.3 | 166.6 | 183.8 | 143.1 | 129.7 |
| **Voles** | 54.3 | 48.1 | 26.7 | 7.0 | 34.0 |
| **Shrews** | 1.4 | 7 | 0.1 | 2.8 | 2.8 |
| **Mustelids** | 0.9 | 0.3 | 0 | 0.3 | 0.4 |

**Table 3.** Terrestrial small mammal species found and their capture rates from deployment of three tunnel types containing trail cameras: the Modified Littlewood tunnel, the Mostela tunnel, and the plastic tunnel. 0* indicates that the species was detected at this tunnel design, but the data were not included in visitation calculations or statistical analysis due to that camera triplet being incomplete and therefore unsuitable for comparison. Rows such as ‘all shrews’ include the counts of the species listed, in addition to individuals which were not identified to species level. Mice were rarely identified to species level and therefore individual species are not listed.

| Scientific name | Common name | Captures in Littlewood tunnel | Captures in Mostela tunnel | Captures in plastic tunnel | Total captures |
| --- | --- | --- | --- | --- | --- |
| *Sorex minutus* | Eurasian pygmy shrew | 13 | 15 | 0 | 28 |
| *Sorex araneus* | Common shrew | 21 | 18 | 4 | 43 |
| *Neomys fodiens* | Eurasian water shrew | 0* | 2 | 0 | 2 |
| **All shrews** |  | **34** | **35** | **4** | **120** |
| *Myodes glareolus* | Bank vole | 373 | 539 | 161 | 1058 |
| *Microtus agrestis* | Short-tailed field vole | 32 | 38 | 26 | 96 |
| **All voles** |  | **503** | **600** | **236** | **1339** |
| **All mice** |  | **2380** | **4,046** | **1693** | **8119** |
| *Rattus norvegicus* | Brown rat | 0* | 0* | 1 | 1 |
| *Sciurus carolinensis* | Eastern grey squirrel | 43 | 23 | 122 | 188 |
| *Mustela nivalis* | Least weasel | 11 | 0* | 1 | 12 |
| *Mustela putorius* | European polecat | 0 | 0* | 0 | - |
| **Lagomorphs** |  | **12** | **0** | **0** | **12** |
| **Total** |  | **3422** | **5316** | **2248** | **11018** |

#### Acoustic Data

The AudioMoths were configured to trigger at frequencies between 20-115kHz, but we found that they were triggered and recorded continuously, despite often no visible noise between this frequency visible on spectrograms. They recorded for an average of 57 hours before memory cards became full or batteries ran out; we found that storage cards (128GB and 64GB) were usually the limiting factor in AudioMoth lifespan.

There were 20 deployments where both acoustic and camera trap data were complete (Table 1), and acoustic recordings were extracted from this subset of data for upload to the BTO Acoustic Pipeline. This subset included deployments from three sites, not including the Black Mountains, and included data from 11 plastic tunnel deployments, and 9 Mostela tunnel deployments. Over the 20 deployments used, the AudioMoths had recorded for a total of 1,178 hours, or 49 days. Files which overlapped in time with a small mammal detection by the cameras were then uploaded to the BTO Acoustic Pipeline; this amounted to 7,128 files (182.5GB).

Of these 7,128 files, 133 five-second terrestrial small mammal records were found representing 4 species: house mouse (12 records), wood mouse (118 records), brown rat (7 records), and common dormouse (8 records). The pipeline also detected 13,828 bat records (12 species), 9,198 bush cricket records (3 species), and 171 moth records (2 species).

The BTO acoustic pipeline assigned a confidence level to each classification, with 89% of classifications being above 0.5 confidence, and 70% of its classifications being above 0.75. Eighteen small mammal species classifications were assigned a confidence level of below 0.50 and we were unable to classify these using a manual inspection, so these records were deemed inconclusive.

### Tunnel performance

#### Data quality

Of complete triplets, 4,204 videos were recorded in the Modified Littlewood tunnels, 3,598 in the plastic tunnels, and 4,162 in the Mostela tunnels. Of these, many videos were non-target triggers – 28.0% in the Modified Littlewood tunnel, 44.7% in the plastic tunnel, and statistically significantly fewer in the Mostela tunnel, at 1.5% (Table 4). Most non-target triggers in the Modified Littlewood tunnel and the plastic tunnel were caused by vegetation moving outside, with very few triggers caused by large mammals passing by.

**Table 4.**
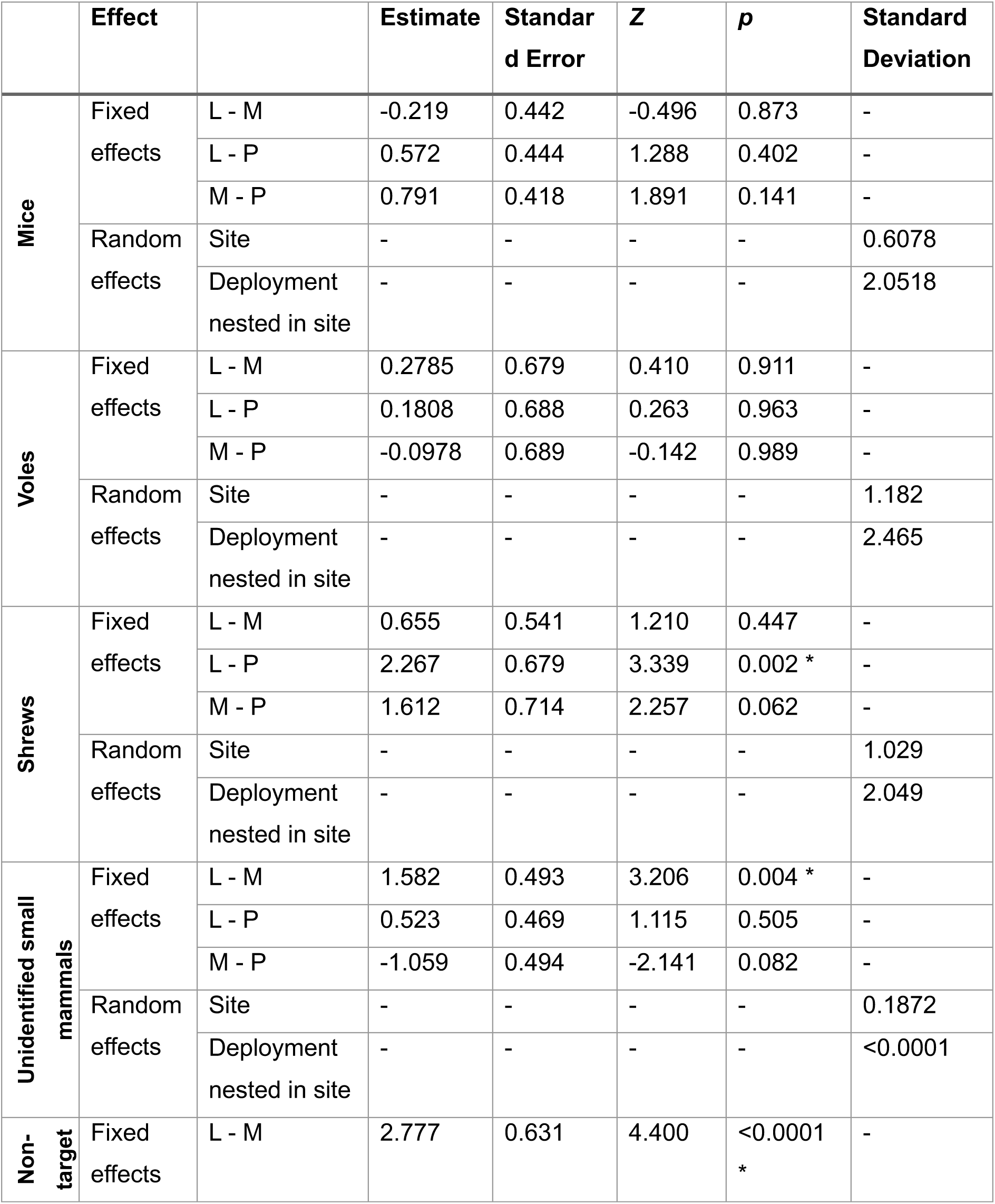
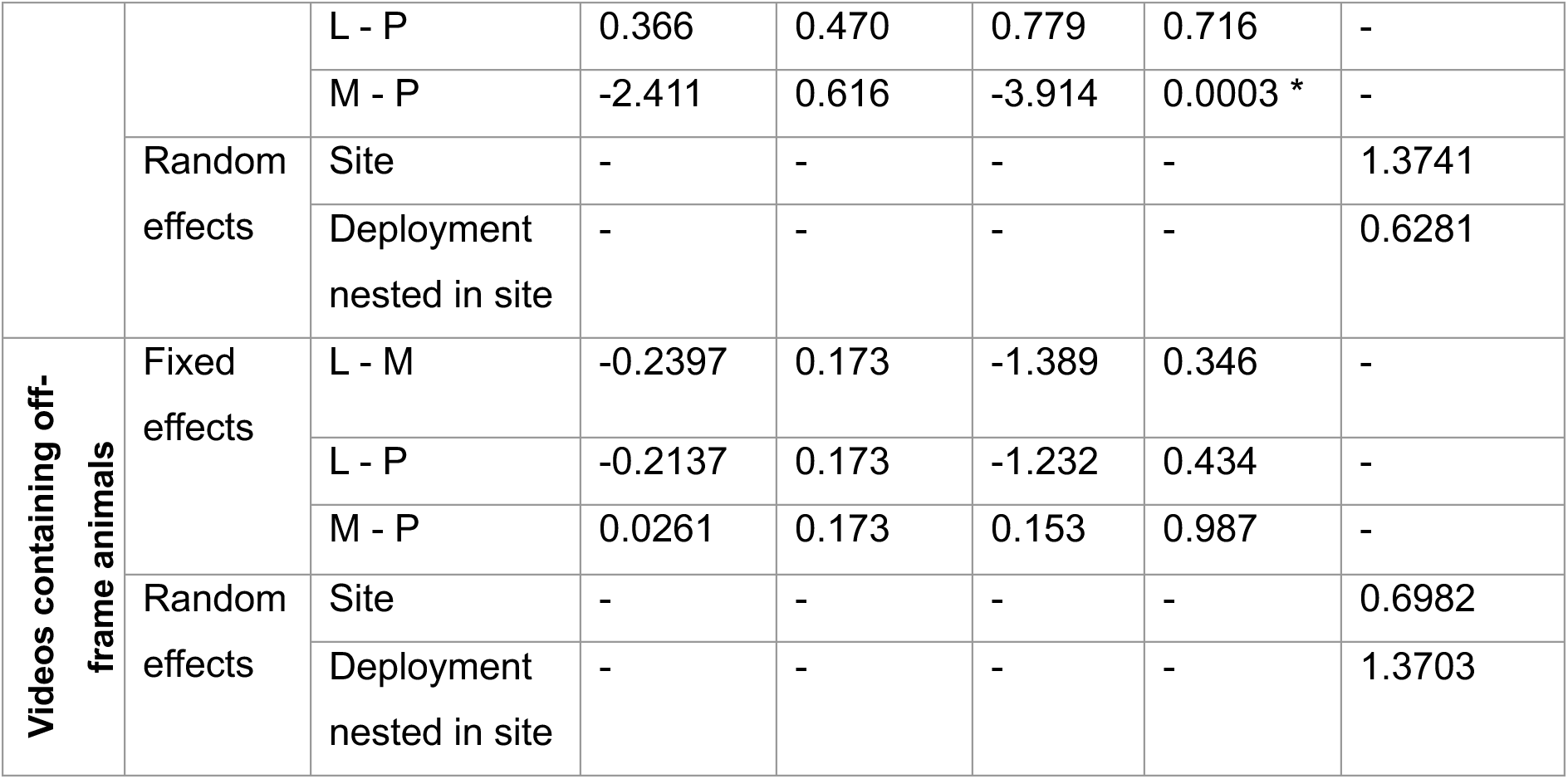
Tukey-adjusted post hoc pairwise comparison statistics of fixed effects, and standard deviations of random effects, from negative binomial generalised linear mixed models comparing count data obtained from camera trapping in three different tunnel designs: the Modified Littlewood tunnel (L), the Mostela tunnel (M), and the plastic tunnel (P). Pairwise comparisons were conducted using estimated marginal means and results are given on a log scale. * highlights significant differences where adjusted p value < 0.05.

Small mammals were unidentifiable into the groupings of vole, shrew, murid, squirrel, or mustelid in 5% of videos in both the Modified Littlewood and plastic tunnel, and 1% of videos in the Mostela tunnel – the GLMM showed that the number of unidentifiable small mammals was significantly lower in the Mostela tunnel compared to Modified Littlewood tunnel (Table 4).

Of small mammal videos, one or more small mammals were off-frame in 27% of videos in the Modified Littlewood tunnel, 29% of videos in the Mostela tunnel, and 51% of videos in the plastic tunnel, however these differences were non-significant (Table 4).

#### Small mammal visitations

Pairwise comparisons of GLMMs showed that there was no significant difference in the number of visitations to each tunnel type for mice nor voles; however, shrews were detected significantly less frequently by the plastic tunnel compared to the Modified Littlewood tunnel (Table 4, Figure 5). There was a non-significant trend towards fewer shrew detections by the plastic tunnel than the Mostela tunnel (*p* = 0.062). Variance among sites was smaller than among deployments nested within sites for mice, voles and shrews (Table 4).

**Figure 5.**
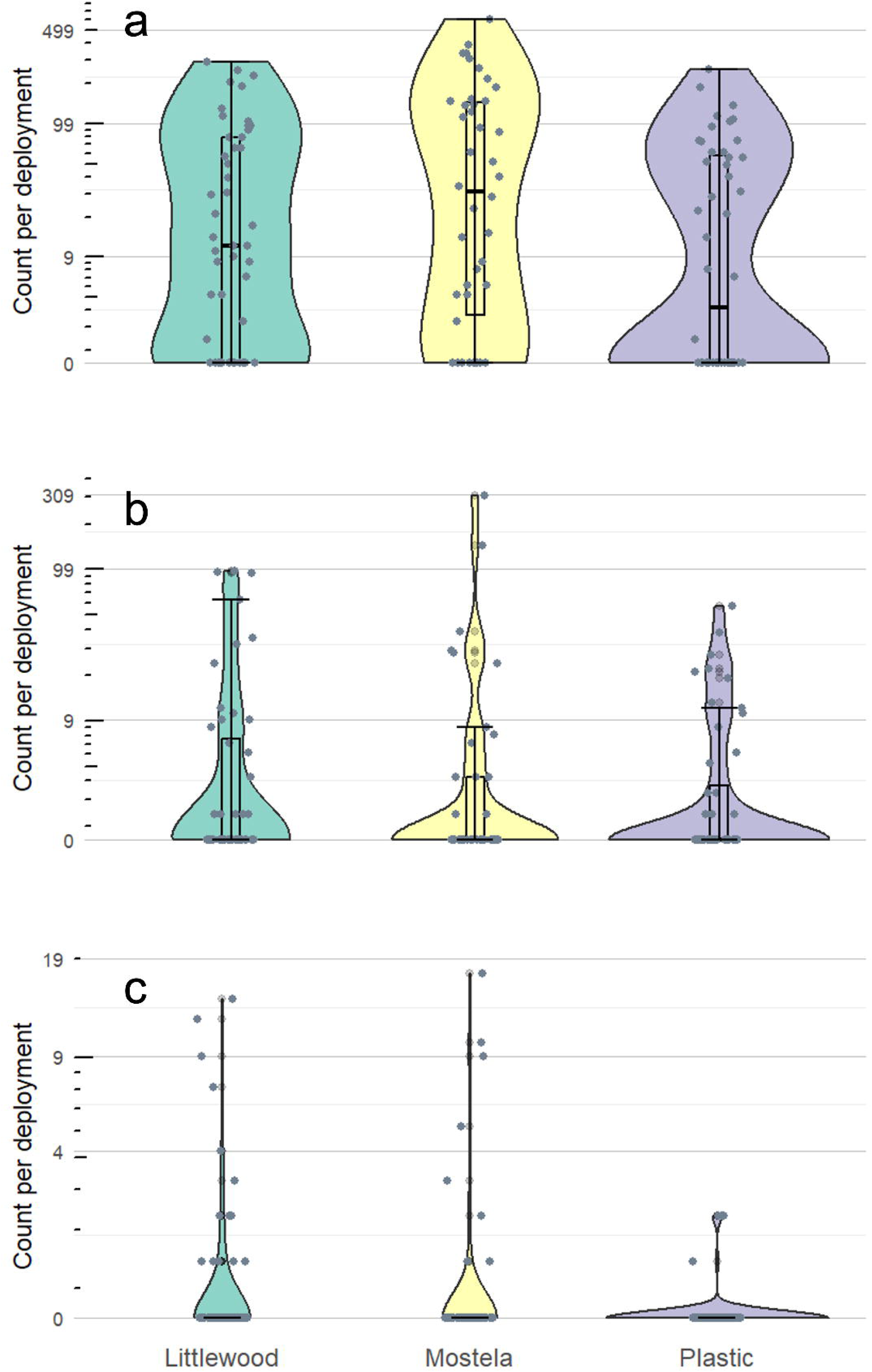
Distribution of count per deployment of terrestrial small mammals detected by trail cameras at three tunnel types in Britain. Boxplots within violin plots show medians and standard deviations.

On average, there were 9 small mammal detections per night in the Modified Littlewood tunnel, 6 per night in the plastic tunnel, and 12 in the Mostela tunnel.

Analysis showed no significant difference in in latency to first detection between tunnel designs for mice (overall average 37.6 hours), voles (overall average 44.8 hours) nor shrews (overall average 98.1 hours) (Table 5, Table 6).

**Table 5.**
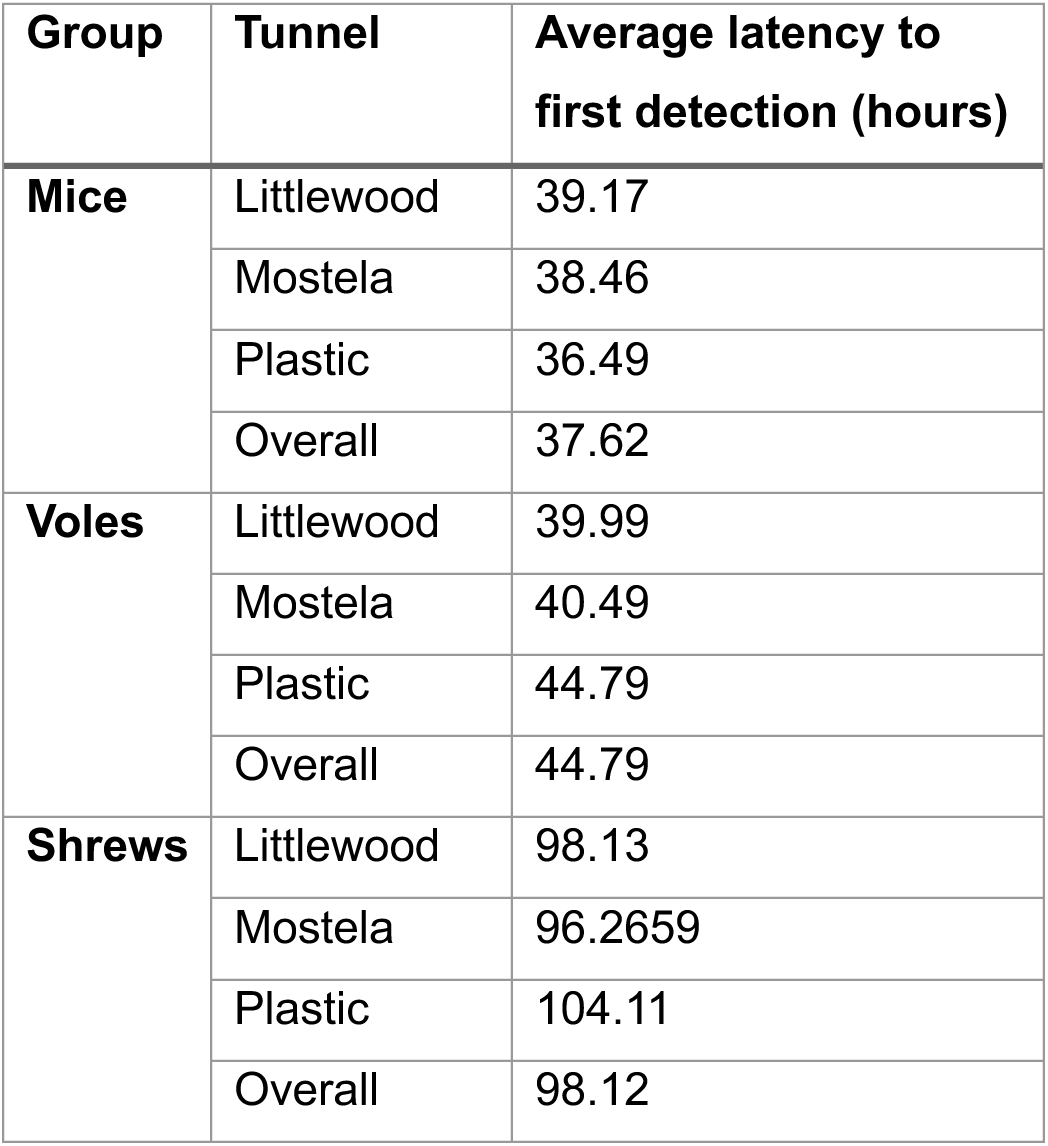
Average latency to first detection of small mammal taxonomic groups in three camera trapping tunnel designs.

**Table 6.**
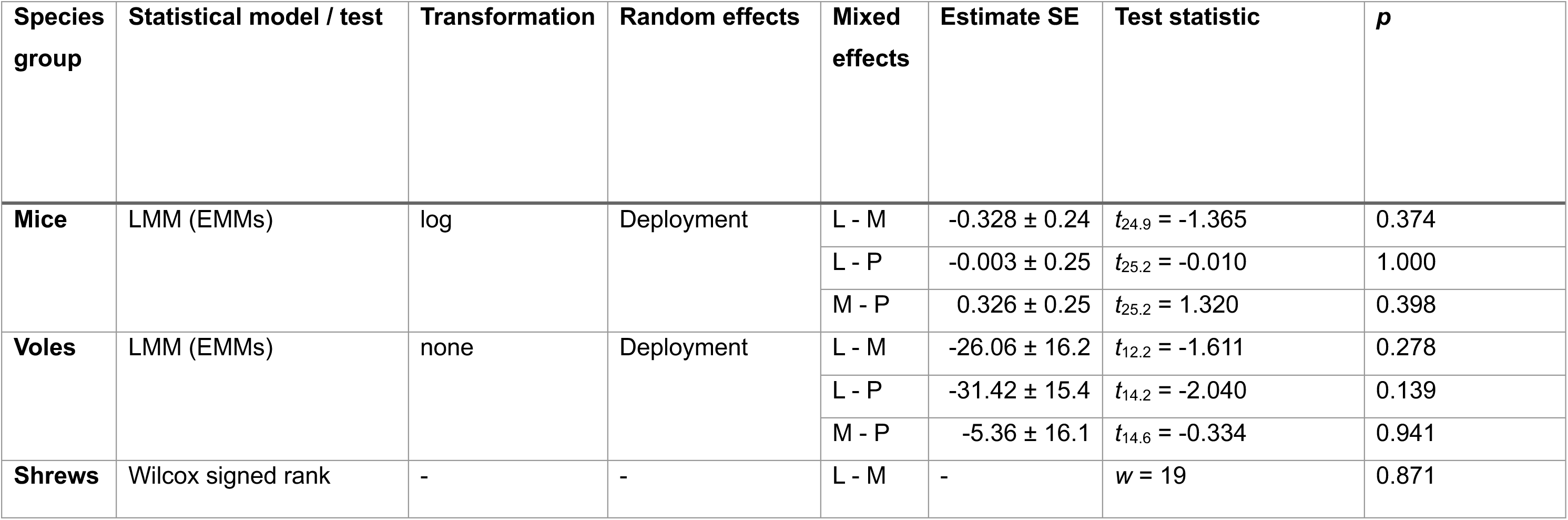
Test statistics from models comparing latency to first detection of mice, voles, and shrews for three camera trap tunnel designs: the Modified Littlewood tunnel (L), the Mostela tunnel (M), and the plastic tunnel (P). Mice and vole statistics are based on linear mixed models (LMMs) with Tukey-adjusted *post hoc* comparisons using estimated marginal means (EMMs).

| Species group | Statistical model / test | Transformation | Random effects | Mixed effects | Estimate SE | Test statistic | *p* |
| --- | --- | --- | --- | --- | --- | --- | --- |
| **Mice** | LMM (EMMs) | log | Deployment | L - M | -0.328 ± 0.24 | $t_{24.9} = -1.365$ | 0.374 |
| | | | | L - P | -0.003 ± 0.25 | $t_{25.2} = -0.010$ | 1.000 |
| | | | | M - P | 0.326 ± 0.25 | $t_{25.2} = 1.320$ | 0.398 |
| **Voles** | LMM (EMMs) | none | Deployment | L - M | -26.06 ± 16.2 | $t_{12.2} = -1.611$ | 0.278 |
| | | | | L - P | -31.42 ± 15.4 | $t_{14.2} = -2.040$ | 0.139 |
| | | | | M - P | -5.36 ± 16.1 | $t_{14.6} = -0.334$ | 0.941 |
| **Shrews** | Wilcox signed rank | - | - | L - M | - | $w = 19$ | 0.871 |

## Passive acoustic monitoring

### PAM and camera trapping

An exact McNemar test showed that there was no significant difference in species group detections by PAM and camera trapping (*χ*^2^1 = 2.77, *p* = 0.096). Of the 59 cases, there were 10 cases of a species being detected by video only, three cases of species being detected by PAM only, and 46 concordant cases. An exact McNemar test on paired vole data (*n* = 14) suggested that camera trapping detected voles on significantly more deployments than did PAM (*χ*^2^1 = 4.17, p = 0.041).

The AudioMoths recorded vocalisations which matched with the species detected by the camera traps in 78 videos, all of which were mice or rats. In all six cases where the recorded vocalisations did not match the species detected by camera trap, common dormouse vocalisations were recorded by the AudioMoth and classified by the BTO, while mice were detected by the camera trap.

### Vocalisation and social behaviours

Exact McNemar tests showed that there was no significant association between vocalisations and multiple individuals being detected by the camera trap, when considering both overall complete audio-video deployments (*χ*^2^1 = 1.50*, n* = 56, *p* = 0.220), and when considering only specific audio-matched video files (*χ*^2^1 = 1.33*, n* = 10, *p* = 0.249).

## Discussion

### Camera trapping

#### Tunnel performance

##### Small mammal visitations

There was no difference in the numbers of mice or voles detected between tunnel types, and no difference in latency to detection among tunnel types for any taxa. We did find that that shrews were detected less frequently by the plastic tunnel compared to the Modified Littlewood tunnel. There was also a trend towards fewer shrew detections by the plastic tunnel than the Mostela tunnel, which may reflect the low sample size. Littlewood *et al*. (2021) detected common and pygmy shrews, but not water shrews, and many studies utilising the Mostela tunnel mention bycatch detections of shrews, including pygmy shrew, common shrew, water shrew, and additional species found outside of Britain (Croose and Carter, 2019; Mos and Hofmeester, 2020; Croose *et al*., 2022, 2025; Park and Lim, 2023; Egloff *et al*., 2025; Granata *et al*., 2026).

Shrews may have been detected more frequently in the Modified Littlewood tunnel compared to the plastic tunnel due to species’ preferences or an improved detection ability. Preferences and detection abilities may be influenced by a combination of factors, including tunnel material, shape, size, colour, number of entrances, and detection area – all of which differed between tunnels. These factors may affect how safe or appealing the tunnel feels to small mammals, or how well the camera performs. Material may impact small mammal visitations to different tunnel types. Material can impact audible sound and ultrasound reflection and propagation (Watkins, Eichelberg and Bilal, 2021), or can emit different volatile organic compounds (VOCs) and therefore smell different: plastics and woods, especially soft woods like those used in this study (Pohleven, Burnard and Kutnar, 2019), emit a range of VOCs (Adamová, Hradecký and Pánek, 2020; Ashraf *et al*., 2023) and plastics have also been shown to absorb VOCs easily (Morris *et al*., 2024), which could deter small mammals if olfactory compounds from previous predator visits are absorbed. Wood used to construct small mammal tunnels, such as the marine ply and pine decking planks used in this study, are likely often treated with preservatives, which may produce additional VOCs.

Fink & Jachowski (2025) concluded that the Littlewood tunnel was better than the Mostela tunnel for small mammal monitoring as it had the shortest latency to detection for mice, opossums, and rabbits, in the US. For *Peromyscus sp*. mice, they found there was a mean latency to detection of 2.6 days in the Littlewood tunnel, and 5.6 days in the Mostela tunnel; we obtained much lower latency to detections for *Apodemus* spp. mice of 39 hours in the Modified Littlewood tunnel, and 38 hours in the Mostela tunnel. In their study, only the Mostela tunnel detected shrews, and only the Littlewood tunnel detected the woodland vole (*Microtus pinetorum*), although this was thought to be due to low population densities. It is possible that small mammal preferences for certain tunnel designs in our study were hidden by the broad classification of small mammals into groupings of ‘vole’, ‘mouse’, and ‘shrew’. Our study had 16 triplet deployments of 7 days, whereas Fink & Jachowski (2025) had 30 triplet deployments of 30 days; some differences may not have been detectable with our smaller sample size.

##### Data quality

Data quality differed between tunnels, with the Mostela tunnel producing the best quality video data – fewer non-target triggers than both other tunnels, and fewer unidentifiable small mammals compared to the Modified Littlewood tunnel.

##### Non-target triggers

The Mostela tunnel produced fewer non-target triggers compared to the Modified Littlewood and plastic tunnel. Non-target triggers accounted for 1.5% of videos in the Mostela tunnel, 28.0% in the Modified Littlewood tunnel, 44.7% in the plastic tunnel.

Non-target triggers were caused by moving vegetation and non-target animals such as large mammals, birds, and invertebrates. Moving vegetation accounted for most of the non-target triggers, with one deployment containing over 1,000 videos thought to be triggered by moving vegetation. The Mostela tunnel likely had fewer non-target triggers than the other two tunnels as cameras did not face the outside and therefore were not triggered by moving vegetation or animals larger than were able to enter the PVC pipe. However, this did mean that the Mostela tunnel was unable to detect small mammals passing by and could only record direct visits to the tunnel. Despite this reduced detection area, the Mostela tunnel performed as well as the Modified Littlewood and plastic tunnel for all small mammal groups. This suggests that the possible advantage of more frequent detections in recording passing-by individuals by the open fronted designs is not worth the additional non-target trigger events, which significantly increased processing time. Any design of small mammal tunnel would likely benefit from the camera not facing the outside.

Some videos which were classified as non-target triggers may have been triggered by a small mammal which then spent the entire video off-frame. On several occasions, no small mammal was visible during the video but noises suggested that something was present and eating off-frame. These were recorded as non-target triggers, as we focused only on the visible data from camera trapping, however it demonstrates that the off-frame area may have increased the number of non-target triggers.

##### Unidentifiable small mammals

Small mammals from videos taken in the Mostela tunnel were more easily identified compared to the other tunnels. Small mammal identification was sometimes not possible because condensation had formed under the camera lens, the lighting was poor, the animal was moving too fast, was already leaving and only partly visible, or because the animal was partly or completely off-frame, or too close to the camera.

The Mostela tunnel may have facilitated small mammal identification due to the profile-view footage obtained as animals enter and leave via the pipe. A profile-view is often useful, especially for voles and shrews where tail-to-body ratio is an important measure of species classification. Several studies have compared species identification rates between profile-view and birds-eye-view camera trap footage, with conflicting results (Smith and Coulson, 2012; Taylor, Goldingay and Lindsay, 2013; Diggins *et al*., 2022); however so far there has been no comparison of species identification rates between profile and head-on views, despite some small mammal camera traps placing the cameras opposite entrances (Littlewood *et al*., 2021; Smaal and van Manen, 2022).

Due to high contrast between the black interior of the plastic tunnel and sunlight from outside, identification was sometimes difficult in the plastic tunnel as only the animal’s silhouette was visible, and only when in front of the entrance. This made it difficult to see differences in colour and tone, and to see body parts which were not directly in front of entrance. This may have increased the number of unidentifiable small mammals in the plastic tunnel.

##### Off-frame individuals

Despite the differences in tunnel shape and size, there was no difference in the number of off-frame individuals between tunnel designs. Small mammals being off-frame meant they could not be identified, and behaviours could not be classified. Any tunnel design could be modified to restrict access to areas beyond the camera’s view and reduce off-frame time. This could be done by addition of funnel shaped barriers, similar to the mesh walls used by Mos & Hofmeester (2020) to separate the camera from visiting small mammals. A barrier preventing small mammals from coming too close to the camera would also facilitate small mammal identification, as small mammals could not be identified when sat directly in front of the camera. Future studies should adapt tunnels to reduce off-frame area to reduce non-target triggers and therefore processing time, improve species classification rates, and obtain more behavioural data.

##### Other considerations

As well as performance, practicality, sustainability, and costs should also be considered. The Mostela tunnel is easily adaptable and can be fitted with a hair tube, tracking tunnel, or measuring tape to facilitate species identification. Due to its size, the Modified Littlewood tunnel could not fit an AudioMoth and was therefore not deployed with one. The plastic tunnels were lighter and did not get heavier when wet, and do not suffer from mould and rot and therefore have a longer lifespan, which further reduces costs. The lifespan of the wooden boxes could be prolonged by attaching wooden batons or feet to the underside to raise it from the ground and reduce rotting. This would make them more cost efficient as the initial costs would be spread over a longer period.

However, the longevity of plastics poses other problems. On several occasions small mammals were observed chewing the sides of the plastic tunnel and the PVC pipe in the Mostela tunnel. Microplastics are of increasing concern to human health and the environment, and have already been found in the bodies of several species terrestrial small mammals, including field voles, wood mice, brown rats, hedgehogs, and lagomorphs (Hornek-Gausterer *et al*., 2021; Alvarez-Andrade *et al*., 2023; Molbert *et al*., 2025; Webber *et al*., 2025), with Thrift *et al*. (2022) finding that 16.5% of faecal samples from British terrestrial small mammals contained microplastics. While the scale of microplastics entering the ecosystem through small mammals chewing on monitoring tunnels is perhaps negligible compared to other sources such as poor waste management (Rohais *et al*., 2024), conservation and research should aim to minimise all negative environmental impacts. Monitoring of threatened species in particular should consider this risk, as target species may already be suffering from reduced reproductive success. Additionally, the purchase of new plastic piping and boxes will contribute to overall plastic waste when retired.

### Acoustic monitoring

#### Camera Trapping *versus* Bioacoustics

During sessions when cameras and AudioMoths were co-deployed, the cameras produced over 3,500 positive small mammal identifications, detecting mice (likely *Apodemus* spp. and house mice), brown rats, common shrews, pygmy shrews, bank voles, and field voles, whereas PAM produced 133 positive small mammal identifications, detecting house mice, wood mice, brown rats, and common dormice. Despite this, the exact McNemar tests suggests that PAM and camera trapping perform equally well for overall small mammal species detection. However, with only 13 discordant pairs, our test had low statistical power. A larger sample size would help confirm this result, and additional data is certainly needed to determine the relative performance of camera trapping and PAM for shrews, mice, rats and dormice, which each had fewer than 4 discordant pairs.

Performing the test on data of individual species rather than species groups would have given the test more statistical power, however detections of individual species were low, and mice could not be identified to species by video. The dataset used for audio-video comparison was much smaller than the camera trapping dataset, due to the reduced active time of the AudioMoths. AudioMoths stopped recording after an average of 57 hours (between 17.6 and 75.8 hours) due to storage filling up or battery failure. This likely limited the number of possible acoustic detections for some species – shrews had an average latency to detection of 98 hours and were only detected by camera traps on two occasions while the AudioMoth was still active.

An exact McNemar test on paired vole data suggested that camera trapping detects voles significantly more frequently than PAM. Vole pups produce isolation calls when alone (Mandelli and Sales, 2004; Szentgyörgyi, Kapusta and Marchlewska-Koj, 2008), adult voles mostly use vocalisations in conspecific communication such as during courtship or aggressive behaviour (Kapusta and Sales, 2009; Kapusta, 2011; Kapusta and Kruczek, 2016). Voles were usually seen alone in this study’s camera trap footage, with under 20 occasions (1% of vole detections) when multiple voles were seen in one video. Additionally, Newson & Pearce (2022) suggest that voles rely more on olfactory cues and do not call as frequently as other small mammals. Therefore, this study may not have detected voles through acoustic analysis as they may vocalise infrequently, perhaps limited to social occasions, which were few in the dataset used for acoustic analysis (two interspecific occasions and two intraspecific occasions).

On several occasions common dormouse calls were recorded by the AudioMoths when a mouse was recorded by the camera trap, suggesting that the AudioMoths can record terrestrial small mammal vocalisations, in addition to bat vocalisations, from outside the tunnels. Camera traps may not have detected dormice as they are arboreal and rarely travel over ground (Bright and Morris, 1991; Bright, 1998), where the camera trap tunnels were deployed. Camera trapping has been successfully used to detect several species of dormouse, however this has usually been done using tree-mounted cameras deployed at heights of between 30 cm and 3 metres above floor level (Di Cerbo and Biancardi, 2013; Mills, Godley and Hodgson, 2016; Mori, Sangiovanni and Corlatti, 2020; Randler and Kalb, 2021), with some as high as 15-31 meters high (Mittelman, Pineda and Balkenhol, 2025). Díaz-Ruiz *et al*. (Díaz-Ruiz *et al*., 2018) also successfully detected dormice through camera trapping but did not specify deployment details. The Dormouse Monitoring System detected garden dormice when deployed 80cm above ground, but not when deployed on the ground (Büchner *et al*., 2022). This suggests that camera traps should be deployed above ground level for dormouse monitoring, and may explain the detection of dormice by the AudioMoths but not by camera traps in the present study. However, a more recent study targeting small mustelids detected edible and garden dormice as by-catch in ground-level Mostela tunnels and external cameras (Granata *et al*., 2026). Further study may therefore be required to determine best practice for dormouse monitoring through direct comparison of arboreal and ground-level monitoring techniques for different species.

PAM required much less processing time than camera trapping, due to the use of an automated classifier. Manually processing the video data took over 60 hours, while the processing the acoustic data took under two hours, including sub-setting the videos and uploading them to the BTO Acoustic Pipeline. While non-invasive techniques such as camera trapping and bioacoustics may require less field time than traditional techniques such as live trapping, they often require more time in data processing. Some simple study alterations could significantly reduce the amount of data collected, such as use of tunnel designs which reduce non target triggers. We recommend using tunnels with entrances perpendicular to the camera to avoid triggers by vegetation and non-target animals. Developments such as WiseEye can reduce non-target triggers by cross-confirming each infra-red trigger using other sensors such as radar (Nazir *et al*., 2017). Additionally, development of automated classifiers such as the BTO Acoustic Pipeline will further reduce processing time, for both video and audio data. Machine learning models can help to both filter out non-target triggers and identify animal species (Norouzzadeh *et al*., 2018; Tabak *et al*., 2019; Yousif *et al*., 2019; Carl *et al*., 2020; Schindler and Steinhage, 2021; Böhner *et al*., 2023). Machine learning classification models can even be built in to camera systems (Whytock *et al*., 2023) and data can transferred across satellite networks (Droissart *et al*., 2021; Whytock *et al*., 2023) to provide instant updates, and reduce field time and ecosystem disturbance. As well as reducing processing time, use of automated classification pipelines will allow non-expert to use bioacoustics and camera trapping, and facilitate citizen science.

Deploying the AudioMoths outside of the tunnels may have resulted in higher quality vocalisation recordings (Stuart Newson, personal communication). Calls may have been distorted by reflection on the insides of the tunnels (Watkins, Eichelberg and Bilal, 2021; Mo *et al*., 2024; Yousefian *et al*., 2025), making the recordings more difficult for the Acoustic Pipeline to classify (Somervuo, Lauha and Lokki, 2023). When analysing audio-matched videos, we assumed that the vocalisations recorded were emitted by the individuals seen in the tunnel on video. However, the recording of dormice while mice were in the tunnel suggests that the AudioMoths were able to record small mammals vocalising outside of the tunnel. Therefore, different individuals may be detected by PAM and camera trapping when both methods detect the same species at the same time.

#### Behavioural information

We found no relationship between vocalisations and number of animals recorded in the traps. Given the low number of small mammal detections by PAM in this study, we were therefore unable to study detailed behavioural associations with calling. Combined use of camera trapping and PAM could allow the study of call types and associated behaviours; however, a much larger sample size would be required than was obtained in this study.

A limitation in the reliability of these results is the difficulty in matching both video and acoustic recording in time. Given that the audiovisual data was obtained through two separate devices and sensors, they may have differing trigger speeds and the time may not be precisely calibrated. We strongly recommend that future studies should find a way to standardise time stamps, such as a clapperboard which is detected by both AudioMoth and camera trap, or for both devices to be linked and synched. This would reduce the time needed for manual analysis.

### General limitations and implications

Data were analysed in species groups because mice were rarely identified to species level, shrews were often not identified to species level, and for shrews and voles, count data for individual shrew species and field vole was too low to analyse separately, with complete video triplets containing 120 total shrew detections, and 96 field vole detections.

Shrew detections were low and varied widely over sites, with 7 detections per camera deployment in the Black Mountains and 0.12 detections per camera at Lower Woods (Table 6). These differences may reflect differences in microhabitat between sites influencing shrew distribution and density. Shrew distribution is known to be influenced by habitat (Shchipanov and Kasatkin, 2023; Balčiauskas *et al*., 2024), vegetation type and structure (Mortelliti and Boitani, 2009; Torre, Díaz and Arrizabalaga, 2014), and water watercourse characteristics such as water depth, current speed, bank incline and bank vegetation (Greenwood, Churchfield and Hickey, 2002). Shrew detections may have been low because little of their preferred microhabitat were sampled. The video data suggests that there were only two shrew visits while AudioMoths were active. This might explain why no shrews were detected by acoustic monitoring in the current study, despite Newson & Pearce (2022) and Vilalta (2024) finding that shrews were the most frequently detected taxon in PAM surveys compared to mice and voles.

Field vole detection also varied widely between sites, with 97 out of 101 total detections being from the Black Mountains, 3 detections at Avon Needs Trees, 1 at Fenswood Farm, and none at Lower Woods. This might also be explained by differences in land use and microhabitats, although study sites contained a variety of habitats, including grassland and field margins, which field voles are associated with (Borowski, 2003; Wheeler, 2008; Balčiauskas *et al*., 2024).

While we deployed 26 camera triplets, only 16 triplets produced complete video data. There were 48 AudioMoth deployments, which produced 20 complete paired audio-visual datasets. The primary cause of datasets being incomplete was the loss of video and audio files during data transfers between recording devices and computers. Transfer errors affected video data from 13 tunnel deployments and acoustic data from 8 tunnel deployments. Overall, transfer errors contributed to 7 incomplete video triplets, after the use of recovery software, and 8 incomplete audio-visual pairings. Additionally, 8 audio-visual pairs were incomplete because the AudioMoths had ceased recording before the first small mammal detection.

## Conclusions

This study aimed to compare the performance of three camera trapping tunnel designs for monitoring terrestrial small mammals. We found that the Modified Littlewood tunnel (Littlewood *et al*., 2021) detected significantly more shrews than the plastic tunnel, and the Mostela tunnel (Croose and Carter, 2019) produced higher quality data than both the Modified Littlewood and plastic tunnel. The Mostela tunnel likely facilitated small mammal identification and reduced non-target triggers through side-facing entrances – other camera trap tunnel designs, including the Littlewood tunnel, could benefit from this. Given that the plastic tunnel did not outperform the other tunnel designs in any aspect (other than affordability, build-time, and ease of transport/cleaning), and poses microplastics concerns, we do not recommend the use of this tunnel, and we encourage plastic tunnel designs and components to be avoided when possible, to reduce negative impacts on our ecosystems, and further make monitoring methods minimally invasive.

We also aimed to evaluate the potential for passive acoustic monitoring compared to camera trapping, a more established technique for surveying small mammals. The small acoustic dataset obtained in this study suggests that PAM may perform as well as camera trapping for overall small mammal species detection, however a much larger sample size is needed to confirm this, and to study behaviours associated with vocalisation. Both camera trapping and passive acoustic monitoring detected species which the other method did not, suggesting that PAM may be useful for some but not all small mammal taxa. Further research is needed to establish the relative performance of bioacoustics for small mammal monitoring compared to other current methods such as live trapping and eDNA surveys.

Camera trapping and PAM can further replace invasive techniques through use alongside methods such as hair sampling, weighing, or a combined approach such as the Small Mammal Monitoring Unit (Bosch *et al*., 2016) and Dormouse Monitoring System (Büchner *et al*., 2022). Through developing our monitoring techniques, we can reduce the use of invasive methods, increase monitoring reliability and accuracy, and improve efficiency by developing automated data processing pipelines. These improvements will benefit species conservation and the study of animal behaviour.

## Appendix 1. Detailed site characteristics

### Fenswood Farm

The site contains a reservoir and stream, and several field edges are kept as wildflower areas. Fenswood Farm is close to a town and city, and surrounding land is also farmland. The hedges and wooded areas of Fenswood Farm are dominated by broadleaf species including oak (*Quercus spp*.), hawthorn (*Crataegus monogyna*), blackthorn (*Prunus spinosa*), hazel (*Corylus avellana)*, field maple (*Acer campestre*), and beech (*Fagus sylvatica*). Previous unpublished surveys have shown that wood mice, house mice, *Rattus* spp., bank voles, grey squirrel, common shrew, brown hare, European rabbit, least weasel, and European polecat. Larger mammals such as European otter, European badger, and roe deer are also present. Annual mean minimum and maximum temperature between 1991-2020 were 7.33 °C and 14.53 °C, and average annual precipitation was 819.01 mm (Filton, ID:676). From May – June 2024, average minimum and maximum daily temperature was 10.6-15.4 °C (Almondsbury, ID: 62122), and average daily rainfall was 1.9 mm (Clifton: Oakfield Road, ID: 9621) (Met Office, 2025).

### Lower Woods

The site is largely Priority Habitat, and consists mostly of broadleaf ancient and semi-natural woodland, with some ancient replanted woodland. Tree species present include oaks (*Quercus petraea, Q. robur*), ash (*Fraxinus excelsior*), hazel, small-leaved lime (*Tilia cordata*), wayfaring tree (*Vibernum lantana*), and wild service tree (*Sorbus torminalis*), with some non-native species including Norway spruce (*Picea abies*). Some areas of hazel are coppiced on a seven-year rotation. Annual mean minimum and maximum temperature between 1991-2020 were 7.33 °C and 14.53 °C, and average annual precipitation was 819.01 mm (Filton, ID:676). Between July and August 2024 when fieldwork took place at Lower Woods, average minimum and maximum daily temperature was 13.1-19.2 °C (Westonbirt, ID: 691) and average daily precipitation was 2 mm (Badminton S WKS, ID: 57110) (Met Office, 2025).

### Great Avon Wood

The site is a forest regeneration project on land that was farmed until 2023, when it was then planted with native broadleaf trees including aspen, alder, hornbeam, pedunculate oak, hawthorn, hazel, field maple and white willow. There was no record of previous small mammal surveys at this site. Annual mean minimum and maximum temperature between 1991-2020 at the nearest weather station were 6.89°C and 14.66°C, and average annual precipitation was 829.92 mm. Between August and September 2024 when this study’s fieldwork took place, average minimum and maximum daily temperature was 10.6-15.4°C (Almondsbury, ID: 62122), and average daily precipitation was 3.6 mm (Bath: Claverton, ID: 56779) (Met Office, 2025).

### Black Mountains

The site, based on a smallholding contains a mixture of apple orchards, fields grazed by Soay sheep, mixed broadleaf woodland, coniferous woodland, and unmaintained regenerating land. The land was grazed intensively by sheep until 2003, after which grazing was reduced and some areas were planted with apple trees or the land was left to regenerate with no interference. No previous small mammal surveys had been performed on site, but least weasel, field vole, and bank vole had been sighted. Predominant species present in hedges and woodlands included oak, hazel, willow, crab apple, blackthorn, hawthorn, alder, elder and bird cherry. Between 1991-2020 the annual mean minimum and maximum temperature at the nearest weather station, Sennybridge 2, was 4.79 °C and 12.19 °C, and average annual rainfall was 1565mm (Met Office, 2025. In June 2024, average daily minimum and maximum temperate was 9.2 °C and 15.3 °C (Libanus, ID: 1249), and average daily precipitation was 0.5 mm (Talgarth W WKS, ID: 56231) (Met Office, 2025).

## Appendix 2. Camera configuration

Configuration details are given in Table A1.

## Appendix 3. Audiomoth container design and configuration

To protect AudioMoths from small mammal interference and water damage, we housed them in small plastic boxes (Figure A1). Slices of foam packaging were placed around the edges of the AudioMoth to ensure a secure fit within each plastic box. A 16 mm hole was drilled above where the microphone would be to allow sound to enter without being altered by the plastic. Middleton *et al*. (2023) suggest that microphones should be deployed within 5 metres of mice and voles to obtain useful acoustic recordings, so we deployed AudioMoths inside the tunnels next to the cameras. Velcro strips were used to attach the AudioMoth containers to the back of the tunnels. AudioMoths were deployed with 128GB storage cards when possible, but 64GB cards were sometimes used. Settings are described in Table A2.

---

**Table A1.** Browning camera settings used for all deployments of terrestrial small mammal monitoring.

| **Camera setting** |  |
| --- | --- |
| Operation mode | Video |
| Video length | 20 seconds |
| Video quality | Ultra |
| Photo delay | 30 seconds |
| Multi shot mode | Off |
| Image data strip | On |
| Motion detection | Long range |
| Trigger speed | Fast |

**Table A2.** AudioMoth configuration settings used for all deployments of terrestrial small mammal monitoring under software version AudioMoth-Firmware-Basic (1.10.0).

| Setting | Configuration |
| --- | --- |
| Sample rate | 250kHz |
| Gain | Medium |
| Enable sleep/record cyclic recording | Sleep 5 seconds, record 55 seconds |
| LED | Enabled |
| Low-voltage cut-off | Enabled |
| Battery level indication | Enabled |
| Schedule | 00:00-24:00 |
| Trigger type | None |
| Filter type | Band pass 20-115kHz |

## FIGURE LEGENDS

**Figure A1.**
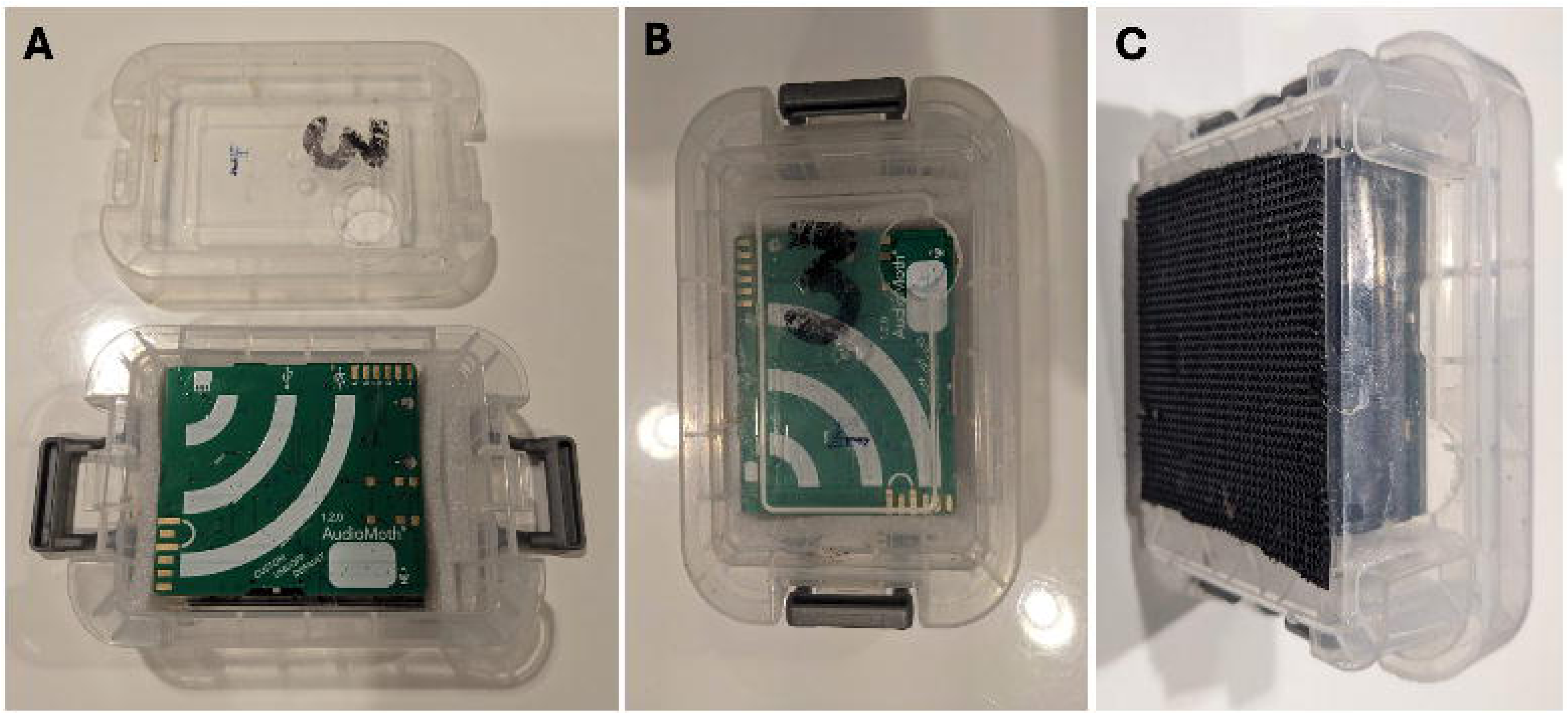
A) AudioMoths were housed in small plastic boxes with slices of packaging foam ensuring a secure fit. B) A 16 mm hole was drilled in the lid above the AudioMoth microphone to allow sound to enter the box unaltered. C) Velcro was stuck to the back of the AudioMoth cases for attachment to the inside of the tunnels.

## DATA ACCESSIBILITY STATEMENT

All data and associated *R* code are available freely at https://doi.org/10.5281/zenodo.18650271

## AUTHOR CONTRIBUTIONS

**LMS:** Conceptualization, Data Curation, Formal Analysis, Investigation, Methodology, Visualization, Writing – Original Draft Preparation, Writing – Review & Editing

**AW:** Conceptualization, Methodology, Resources, Supervision, Writing – Review & Editing

**SAR:** Conceptualization, Methodology, Supervision, Writing – Review & Editing

## ACKNOWLEDGMENTS

We would like to thank Stuart Newson for his input in designing our passive acoustic monitoring methodology. We are also grateful to Avon Needs Trees, Gloucestershire Wildlife Trust, Fenswood Farm (University of Bristol), and Menna Angharad and Jeremy Stuff from the Black Mountains smallholding for granting us field permissions and providing assistance.

